# A cereblon (CRBN) molecular glue degrader of NIMA-related kinase 7 (NEK7) reveals a context-dependent role in NLRP3 inflammasome activation

**DOI:** 10.1101/2024.11.06.622079

**Authors:** Aude Sylvain, Natacha Stoehr, Fupeng Ma, Artiom Cernijenko, Martin Schröder, Maryam Khoshouei, Melanie Vogelsanger, Michel Schoenboerner, Ashley Burke, Rao Pasupuleti, Jonathan Solomon, Joshiawa Paulk, Lei Xu, Janet Dawson, Damien Begue, Peggy Lefeuvre, Erik Ahrne, Andreas Hofmann, Callum J. Dickson, Philip Arabin, Alfred Zimmerlin, Michael Kiffe, Mirjam Froehlicher, Theresa Boersig, Azeddine Elhajouji, Murielle Brichet, Sue Menon, Shanming Liu, Matthias Mueller, Charles Wartchow, James Lin, Yamel Cardona Gloria, Sabine Dickhöfer, Alexander N. R. Weber, Tatjana Welzel, Jasmin Kuemmerle-Deschner, Christopher J. Farady, Robert Pulz, Frederic Bornancin, Dennis L. Buckley, Zuni I. Bassi

## Abstract

The NLRP3 (NACHT-, leucine-rich repeat [LRR]- and pyrin domain [PYD]- containing protein 3) inflammasome is a cytoplasmic signaling complex that promotes inflammation in response to signals from infection and cellular damage. Increased activation of the NLRP3 inflammasome is linked to numerous diseases including gout, osteoarthritis, cardiovascular disease, neuroinflammatory diseases and cancer, which has prompted the search for therapeutics that can inhibit the NLRP3 pathway.

Recent work suggested that NIMA-related kinase 7 (NEK7) may be required for proper assembly and activation of the NLRP3 inflammasome independent of its kinase activity, and that hence reduction of NEK7 protein levels may block NLRP3 activation. Since NEK7 contains a glycine beta hairpin loop structural degron found in many targets of cereblon (CRBN) molecular glue degraders, we used this strategy to degrade NEK7 and inhibit the NLRP3 inflammasome. We identified NK7-902, a CRBN glue degrader of NEK7. NK7-902 potently and selectively degraded NEK7 in human primary monocytes, peripheral blood mononuclear cells (PBMCs) and whole blood. Unexpectedly, full NEK7 degradation led to only partial blockade of NLRP3-dependent interleukin-1β (IL-1β) release in these cells under different stimulation conditions, with the extent of IL-1β inhibition varying greatly across donors.

Unlike most CRBN glue degraders, NK7-902 degraded NEK7 in mouse cells and efficiently inhibited NLRP3-dependent IL-1β release in a mouse cryopyrin-associated syndrome (CAPS) model. By contrast, oral administration of NK7-902 in non-human primates led to profound and long lasting NEK7 degradation but only transiently blocked IL-1β production in blood.

Collectively, our data suggest that NEK7 is involved but may not be strictly required for NLRP3 activation in primates and humans.

**Highlights:**

- Identification of NK7-902, a CRBN molecular glue degrader of NEK7
- NK7-902 fully degrades NEK7 in human primary monocytes and whole blood but only partially inhibits NLRP3-dependent IL-1β production
- Unlike most CRBN glue degraders, NK7-902 shows activity in murine systems
- In vivo oral administration of NK7-902 in non-human primates leads to profound and long-lasting NEK7 degradation but only transiently blocks NLRP3 inflammasome activation

## Introduction

The NLRP3 (NACHT-, leucine-rich repeat [LRR]- and pyrin domain [PYD]- containing protein 3) inflammasome is a cytoplasmic signaling complex that assembles in response to exogenous microbial infections and danger signals such as pore-forming toxins, adenosine triphosphate (ATP), cholesterol crystals and protein aggregates. Through a common activation mechanism associated with cellular K^+^ efflux, these diverse danger signals trigger NLRP3 inflammasome activation, leading to an inflammatory response that involves the production of the pro-inflammatory cytokines interleukin 1β (IL-1β) and IL-18, as well as gasdermin-D mediated pyroptotic cell death ^1^.

Formation of the NLRP3 inflammasome is a tightly regulated process that occurs in two steps ^2^. The first, known as priming/licensing, involves activation of pattern recognition receptors (PRR) such as Toll-like receptor 4 (TLR) by components of microorganisms (e.g. lipopolysaccharides), leading to NF-κB activation. NF-κB-dependent ‘transcriptional priming’ is required to increase expression of the inflammasome components NLRP3, caspase-1 and pro-IL-1β. TLR signaling also triggers post-translational modifications of NLRP3 that stabilize an auto-suppressed inactive form and assist NLRP3 maturation to a signal-competent state. The second step, activation, results in the formation of a disk-shaped NLRP3 oligomer, which recruits the adaptor protein ASC (apoptosis-associated, speck-like protein containing a caspase recruitment domain [CARD]) and the protease caspase-1, leading to caspase dimerization and activation. Activated caspase-1 cleaves pro-IL-1β, pro-IL-18 and the pore-forming protein gasdermin D to produce their active, mature forms and promote inflammation ^3^.

Gain-of-function mutations of the NLRP3 gene cause a group of rare inherited autoinflammatory diseases, collectively known as cryopyrin-associated periodic syndromes (CAPS) ^4^. In addition, NLRP3 hyperactivation has been linked to a variety of conditions, including gout, osteoarthritis, cardiovascular disease, neuroinflammatory diseases and cancer, making this pathway an attractive therapeutic target to treat these diseases ^1^.

NIMA-related kinase 7 (NEK7) was first described to have a role in mitosis; it localizes to the centrosome and regulates mitotic spindle formation and microtubule dynamics as part of a signaling cascade involving two other NIMA-family members, NEK9 and NEK6 ^5–11^. Subsequent studies in murine cells also identified NEK7 as a critical cofactor for NLRP3 activation ^12–14^. The functions of NEK7 in mitosis and in the NLRP3 inflammasome were suggested to be mutually exclusive to prevent inflammasome activation during cell division ^14^. A cryo-electron microscopy (cryo-EM) structure of an engineered NEK7 dimer in complex with inactive human NLRP3, showed an interaction between the C-lobe of the NEK7 kinase domain and the LRR and NACHT domains of NLRP3, suggesting that NEK7 may function as a bridge connecting adjacent NLRP3 molecules via their LRR domains ^15^. This function of NEK7 was independent of its catalytic activity and appeared to be specific to the NLRP3 inflammasome ^12,14^, although a recent report suggested that NEK7 may also be involved in NLRP1 activation ^16^.

Collectively, a body of evidence supports a role for NEK7 in NLRP3 inflammasome function ^12–15,17–21^. However, the involvement of NEK7 has remained incompletely understood and has been challenged by recent studies. For instance, an IKKβ-mediated pathway was shown to enable NLRP3 activation independently of NEK7 in human iPS-derived macrophages and human myeloid cell lines ^22^. Similarly, NLRP3 activation in HEK293 cells co-expressing ASC and NLRP3 occurs in a NEK7-independent manner ^23^. Furthermore, structural analysis of the active, disc-shaped NLRP3 oligomer in complex with adenosine 5’-O-(3-thio)triphosphate, NEK7, and ASC revealed that the C-terminal LRR domain of NLRP3 bound to NEK7 extends away from the center of the active oligomer and does not directly participate in disc interfaces ^24^, suggesting that NEK7 may play a less critical role in NLRP3 activation than previously thought. More recently, two paths were described accounting for canonical NLRP3 activation: one pathway relying on the assembly of NLRP3 oligomeric cages (decameric in human, dodecameric in mouse) ^21,25^ and their transit to the microtubule organization center (MTOC) via TGN38+ vesicles, and another cytosolic pathway which does not rely on NLRP3 cages or TGN/MTOC association ^25^. This novel cytosolic, cage-independent pathway is likely NEK7-independent; in fact, a similarly cytosolic but C-terminally truncated form of NLRP3 was inflammasome-competent despite being unable to interact with NEK7 ^26^. Thus, in human cells, the requirement for NEK7 needs further investigation, also to explore whether NEK7 could be a therapeutic target for restricting NLRP3-mediated inflammation.

Here, we describe the identification and characterization of a cereblon (CRBN)-molecular glue degrader of NEK7, NK7-902. We show that NK7-902 potently and selectively degraded NEK7 in human primary monocytes, peripheral blood mononuclear cells (PBMCs) and whole blood *in vitro*. Unexpectedly, we found that NK7-902-mediated NEK7 degradation inhibited IL-1β release to varying extents depending on the donor and stimulation conditions, indicating that the NLRP3 pathway can be, at least in part, activated in a NEK7-independent manner in human primary cells *in vitro*. Unlike other CRBN glue degraders, NK7- 902 could effectively degrade NEK7 in mouse and inhibit NLRP3-dependent IL-1β release in a mouse CAPS model to a similar extent as achieved with a direct NLRP3 inhibitor. Finally, we demonstrate profound and long-lasting NEK7 degradation upon oral administration of NK7-902 *in vivo* in non-human primates (NHP) but also show that this leads to only transient and partial IL-1β inhibition in *ex vivo* stimulated blood in this species.

Thus, our findings concur and add substantially to the accumulating evidence that, in primates and humans, NEK7 is involved but not strictly required for NLRP3 activation.

## Results

### Discovery of NK7-902, a CRBN molecular glue degrader of NEK7

Targeting NEK7 via induced degradation is an attractive approach because the role of NEK7 in NLRP3 activation is known to be independent of its kinase activity. It was previously demonstrated that CRBN glue degraders, such as pomalidomide, bind their targets through a structural degron known as a g-loop or glycine containing β-hairpin ^27^. Analysis of NEK7 revealed the presence of a β-hairpin structural degron (NEK7 residues 51-60) with high similarity to the β-hairpin of Casein Kinase 1α (CK1α) (RMSD 0.25 Å), suggesting that NEK7 might be targetable by a CRBN glue degrader. We screened 2,500 CRBN-binding molecules for their ability to recruit NEK7 to CRBN using a cell-based split enzyme recruitment assay, analogous to those previously used for IKAROS Zinc Finger 1 (IKZF1) and IKZF2 ^28^. One compound, NK7-288 not only recruited NEK7 to CRBN but also induced weak NEK7 degradation in a high-throughput degradation assay (Figure 1A-C). The replacement of the isoindolinone of NK7-288 with a phthalimide led to NK7-147, which had increased potency in the high throughput NEK7 degradation assay (Figure 1A-C). Further elaboration of the tail moiety led to NK7-902, which had comparable degradation in our high-throughput assay and slightly improved selectivity against IKZF2 degradation (Figure 1A-C and Supplementary Figure 1A).

**Figure 1.**
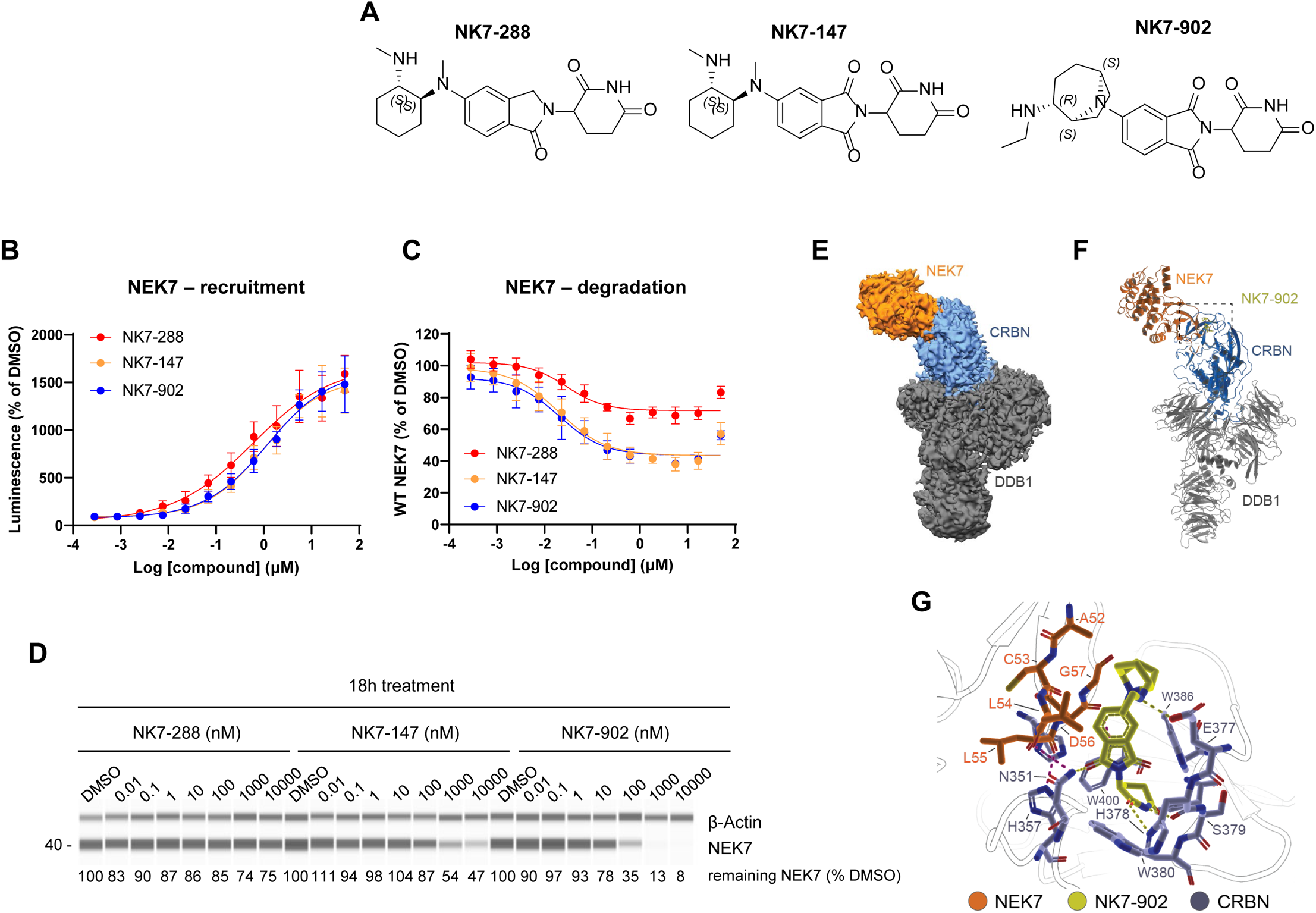
Identification of NK7-902, a CRBN glue degrader of NEK7. (A) Structure of NK7-288, NK7- 147 and NK7-902. (B) Cell-based NanoBiT recruitment assay. 293T cells were transiently transfected with plasmids encoding for NEK7 and CRBN tagged with SmBiT and LgBiT, respectively, and then incubated with increasing concentrations of NK7-288, NK7-147 or NK7-902 for 2h in the presence of the NAE1 inhibitor MLN4924 as described in the methods. Data represent mean ± SD (n=8) (C). ProLabel NEK7 degradation assay. 293T cells were transiently transfected with a plasmid encoding for ProLabel-tagged NEK7 and then incubated with increasing concentrations of NK7-288, NK7-147 or NK7-902 for 18h. Data represent mean ± SD (n=4). (D) THP1 cells were incubated with increasing concentrations of NK7-288, NK7-147 and NK7-902 for 18h and then lysed to measure NEK7 and β-actin levels by WES capillary electrophoresis immunoanalysis. For each sample, the NEK7 signal was normalized over the corresponding β-actin signal. Percentage of remaining NEK7 compared to the DMSO control was calculated. A representative experiment out of two independent repeats is shown. (E) Refined EM-map of the ternary complex of CRBN-DDB1, NEK7 and NK7-902 (pdb ID: 9H59). EM-map is color-coded by the corresponding modeled proteins. (F) Model of the ternary complex of CRBN-DDB1, NEK7 and NK7-902. The dashed box indicates the area highlighted in sub-panel G. (G) Modeled interactions of the ternary complex of CRBN-DDB1, NEK7 and NK7-902. Potential hydrogen bonds between NEK7 and CRBN are highlighted by magenta dashes while NK7-902 interactions to CRBN are displayed as yellow dashes.

To confirm that the compounds were acting on the β-hairpin of NEK7, we generated mutations in the crucial glycine of the β-hairpin and tested the mutants in the CRBN recruitment assay. The G57N mutant abrogated both NEK7 recruitment to CRBN as well as NEK7 degradation, strongly implicating the β- hairpin of NEK7 as the degron (Supplementary Figure 1B-C). The lack of degradation observed in CRBN knockout cells further supported the fact that the compounds were CRBN-based glue degraders (Supplementary Figure 1D). Interestingly, a NEK7 G57A mutant was still recruited to CRBN suggesting that, in the case of NEK7, the alanine residue, although larger than the glycine, still allowed β-hairpin formation and interaction with CRBN (Supplementary Figure 1E). Although NK7-147 and NK7-902 had comparable activity in our high throughput degradation assay (Figure 1C), NK7-902 was able to induce complete degradation of endogenous NEK7 in THP1 cells at a concentration of 1 µM, whereas NK7-147 led to only partial degradation (Figure 1D). In addition, although the cellular recruitment assay was unable to distinguish between the three degraders, NK7-902 achieved stronger recruitment than NK7-147 using recombinant proteins in Time-Resolved Fluorescence Resonance Energy Transfer (TR-FRET) and surface plasmon resonance (SPR) assays (Supplementary Figure 1F and 1G), potentially explaining its improved degradation activity on endogenous NEK7 in THP1 cells.

In order to further validate the β-hairpin-mediated interaction of NEK7 with CRBN in the presence of NK7- 902, we used cryo-EM to obtain a structural model of the ternary complex. The resulting, refined EM map of 3.38 Å resolution (Figure 1E, Supplementary Figure 2A-C) allowed modeling the ternary complex of NEK7, NK7-902 and CRBN-DDB1 (Figure 1F). Despite flexibility of CRBN and NEK7 and the resulting loss of local high resolution (Supplementary Figure 2D), the point mutation studies as well as analyses of previously solved high-resolution structures of CRBN- β-hairpin interactions ^27,29–31^ guided building the interaction model. We observed sufficient density of NK7-902 in the tryptophan pocket as well as the β- hairpin of NEK7 to place both parts (Supplementary figure 2E). NK7-902 showed interactions via the hydrogen bonds with H378 and N351 (Figure 1G). Additionally, the secondary amine interacted with the side chain of E377. NEK7 β-hairpin residues interacted via H-bonds with CRBN and NK7-902 providing shielding effects as previously observed ^32^. The hydrophobic interactions of the bridged ring system of NK7-902 with G57 also gave insights for the preserved recruitment of the G57A mutant, potentially still allowing a similar compound binding mode, while a bulkier residue in the G57N mutant would prohibit NEK7 β-hairpin interactions with NK7-902.

### NK7-902 degrades NEK7 and inhibits the NLRP3 inflammasome in human primary monocytes

We next characterized the ability of NK7-902 to degrade NEK7 in human primary monocytes. NK7-902 induced concentration-dependent NEK7 degradation in these cells, with a DC_50_ of 0.2 nM and D_max_ > 95% (Figure 2A), exhibiting a higher degradation efficacy than in THP-1 cells. NEK7 degradation upon treatment with NK7-902 was time-dependent, with 80% NEK7 degradation achieved within 1 hour (h) after the addition of the compound and maximum degradation seen after 24h (Figure 2B).

**Figure 2.**
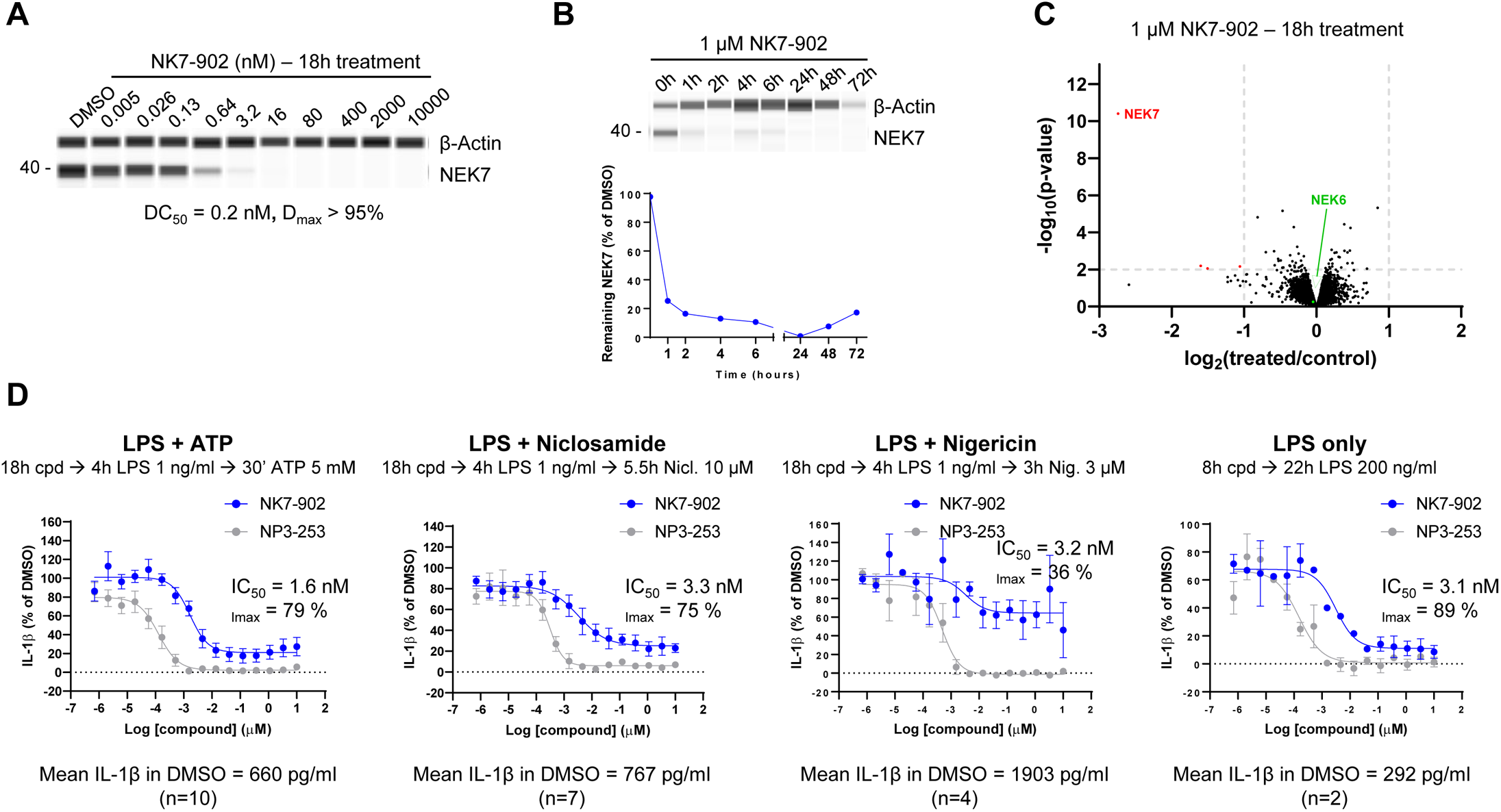
NK7-902 degrades NEK7 and inhibits the NLRP3 inflammasome in human primary monocytes. (A) Human primary monocytes were treated with increasing concentrations of NK7-902 for 18h and then lysed to measure NEK7 and β-actin levels by WES capillary electrophoresis immunoanalysis. A representative experiment out of three independent repeats is shown (see also Supplementary Figure 2A). (B) Human primary monocytes were incubated with 1 µM NK7-902 for the indicated times, cells were lysed and NEK7 and β-actin levels were measured by WES capillary electrophoresis immunoanalysis. For each sample, the NEK7 signal was normalized over the corresponding β-actin signal. Percentage of remaining NEK7 compared to the 0h time-point was calculated and plotted. A representative experiment out of two independent repeats is shown. (C) Human primary monocytes were incubated with 1 µM NK7-902 for 18h and then analyzed by LC-MS/MS to identify differentially modulated proteins. Proteins in the top left quadrant were significantly downregulated (see also Supplementary Table 1). (D) Human primary monocytes were pre-incubated with increasing concentrations of NK7-902 or the NLRP3 inhibitor NP3-253 and then stimulated as indicated at the top of each graph. IL-1β levels in the supernatant were measured by HTRF. The mean IL-1β concentration (pg/mL) in the DMSO control for each stimulation condition is indicated below each graph. IC_50_ and I_max_ values for NK7-902 in the different conditions are shown. Each stimulation condition was tested in monocytes from the indicated number of independent human donors, data represent mean ± SEM (see also Supplementary Figure 2D).

The selectivity of NK7-902 was assessed by mass-spectrometry based proteomic analysis of human primary monocytes following an 18h treatment with 1 µM NK7-902. In this cell type, NEK7 was the most significantly down-regulated protein across all quantified proteins (Figure 2C), highlighting the high selectivity of NK7-902. NEK6, the protein most homologous to NEK7 (86% identical in their catalytic domain) does not contain a β-hairpin and was not degraded by NK7-902, confirming the importance of this structural motif for successful CRBN-mediated degradation (Figure 2C). Expression proteomics analysis of NK7-902 was also done in human induced pluripotent stem (iPS) cells which express Sal-like protein 4 (SALL4), a zinc-finger transcription factor only weakly expressed in monocytes, whose degradation has been linked to IMiD-induced teratogenicity ^33,34^. NEK7 was the most strongly and significantly decreased protein in iPS cells and SALL4 protein levels were detected but unchanged after treatment with NK7-902 (Supplementary Figure 3B). Interestingly, we observed a significant downregulation of SALL3, another member of the SALL gene family (Supplementary Figure 3B). FLT3 interacting zinc finger 1 (FIZ1) and the zinc finger transcription factor IKZF4 were also significantly down-regulated by NK7-902 in human iPS cells (Supplementary Figure 3B).

We next evaluated whether treatment of primary immune cells with NK7-902 can lead to functional inhibition of the NLRP3 inflammasome. Human monocytes were pre-treated with NK7-902 for 18h, then the NLRP3 inflammasome was primed by 4h treatment with lipopolysaccharide (LPS) and activated with either exogenous ATP, the mitochondrial uncoupler niclosamide, or the pore-forming toxin nigericin. Inflammasome activity was determined by measuring NLRP3-dependent IL-1β release. A selective NLRP3 inhibitor, NP3-253 (see Materials and Methods section), was included in all experiments as a comparator. NK7-902 potently inhibited IL-1β secretion in cells activated with either ATP or niclosamide, reaching on average a maximum inhibition of 79% and 75%, respectively, compared to full inhibition seen with the NLRP3 inhibitor NP3-253 (Figure 2D). Of note, the extent of IL-1β inhibition with NK7-902 varied greatly across different donors in these conditions, with some showing full blockade of IL-1β release and some displaying a very limited effect (Supplementary Figure 3C). In addition, only 36% IL-1β blockade was achieved with NK7-902 in monocytes activated with nigericin, with all donors tested showing incomplete inhibition (Supplementary Figure 3C). We also tested whether NK7-902 could inhibit IL-1β release upon activation of NLRP3 via the alternative inflammasome pathway. This pathway exists in human, not rodents, and is induced through stimulation with LPS only, involves caspase-8 activation and does not induce apoptosis-inducing speck-like protein containing CARD (ASC-speck) formation or pyroptosis ^35^. NK7-902 blocked NLRP3-dependent IL-1β release by 89% in these conditions, compared to full inhibition with NP3-253 (Figure 2D). Together, these data suggest that NK7-902 mediated NEK7 degradation can block both canonical and alternative NLRP3 pathway activation *in vitro* but that the extent of IL-1β inhibition is donor- and stimulus-dependent.

### NK7-902 partially inhibits NLRP3-dependent IL-1β release in human whole blood

To understand the consequences of NEK7 degradation in a mixed immune cell population, the effect of NK7-902 on isolated human peripheral blood mononuclear cells (PBMCs) was assessed. NK7-902 potently degraded NEK7 in PBMCs following an 18h incubation (Figure 3A). Similar to what we had observed in monocytes, NK7-902 potently inhibited NLRP3-dependent IL-1β release upon stimulation with LPS plus ATP but the effect was incomplete (71% maximum inhibition) when compared to the NLRP3 inhibitor NP3-253 (Figure 3B).

**Figure 3.**
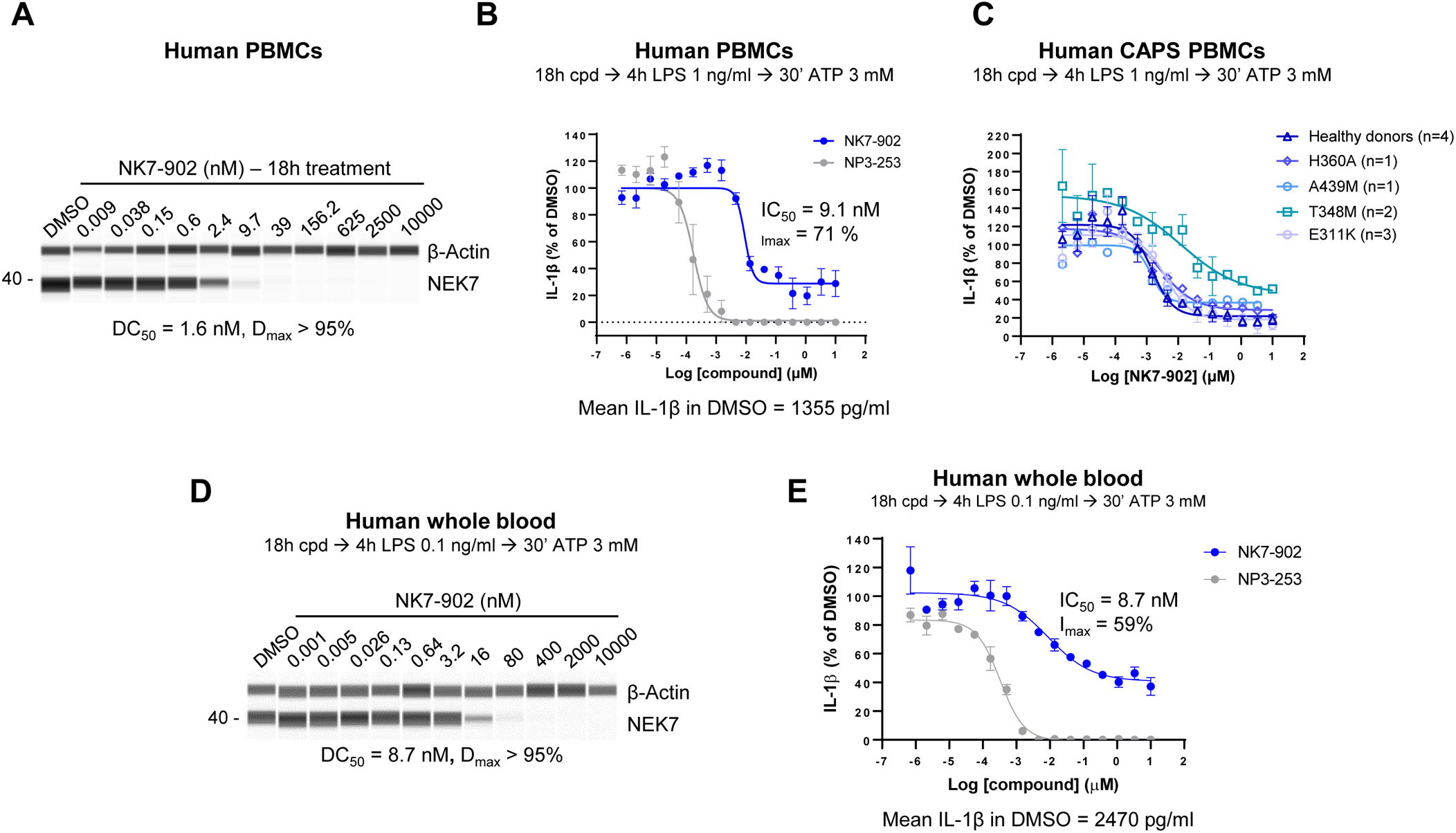
NK7-902 partially inhibits NLRP3-dependent IL-1β release in human PMBCs and whole blood. (A) Human PBMCs were treated with increasing concentrations of NK7-902 for 18h and then lysed to measure NEK7 and β-actin levels by WES capillary electrophoresis immunoanalysis. A representative experiment out of three independent repeats is shown (see also Supplementary Figure 3A). (B) Human PBMCs were pre-incubated with increasing concentrations of either NK7-902 or NP3-253 for 18h and then stimulated with LPS and ATP as indicated at the top of the graph. IL-1β levels in the supernatant were measured by HTRF. The mean IL-1β concentration (pg/mL) in the DMSO control is indicated below the graph. (C) PBMCs from healthy volunteers and CAPS patients carrying different NLRP3 mutations were pre-incubated with increasing concentrations of NK7-902 for 18h and then stimulated with LPS and ATP as indicated. IL-1β levels in the supernatant were measured by HTRF. (D) Human whole blood was diluted 1:2 in medium and incubated with increasing concentrations of NK7-902 for 18h prior to stimulation with LPS followed by ATP. Red blood cells were lysed and NEK7 and β-actin levels were measured in white blood cell protein extracts by WES capillary electrophoresis immunoanalysis. (E) Human whole blood was diluted 1:2 in medium and incubated with increasing concentrations of either NK7-902 or NP3-253 for 18h prior to stimulation with LPS and ATP. IL-1β levels in the plasma fraction were measured by HTRF. The mean IL-1β concentration (pg/mL) in the DMSO control is indicated below the graph. In (B), (C) and (D) data represent mean ± SEM.

CAPS is a group of rare inherited autoinflammatory diseases caused by gain-of-function mutations in the NLRP3 gene, leading to hyperactivation of the NLRP3 inflammasome ^4^. To investigate whether and to which extent these mutation-sensitized NLRP3 proteins may rely on NEK7 for building an inflammasome, PBMCs from healthy donors or from CAPS patients carrying different activating NLRP3 mutations were treated with NK7-902 for 18h and then stimulated to induce IL-1β release. Although CAPS NLRP3 mutants solely require a priming signal (*e.g.,* LPS) to assemble into active inflammasomes, an LPS plus ATP stimulation was chosen for both healthy and CAPS PBMCs to enable a proper comparison between wild-type and mutant forms. The extent and potency of NEK7 degradation mediated by NK7-902 was comparable in PBMCs from healthy donors and CAPS patients (Supplementary Figure 4B). Activation of NLRP3 inflammasomes carrying H360A, A439M and E311K mutations was efficiently blocked by NK7- 902 treatment (Figure 3C). In PBMCs with NLRP3 carrying the T348M CAPS mutation, treatment with NK7-902 only partially reduced IL-1β secretion (Figure 3C), which might suggest a reduced dependency on NEK7 in the presence of this mutation.

Next, we studied the effect of NK7-902 on inflammasome activation in human whole blood, a matrix commonly used to mimic the environment that an oral drug will encounter *in vivo*. NK7-902 effectively degraded NEK7 in human whole blood (Figure 3D) but only partially inhibited NLRP3-dependent IL-1β release in this matrix (59% maximum inhibition) as opposed to NP3-253, which fully blocked production of IL-1β (Figure 3E). Because neutrophils can make up to 75% white blood cells and are important contributors of IL-1β in addition to monocytes, we tested whether NK7-902 could degrade NEK7 and inhibit IL-1β release in neutrophils. NK7-902 induced profound and concentration dependent degradation of NEK7 in human neutrophils and inhibited IL-1β induced by LPS plus ATP stimulation to a similar extent as observed with the NLRP3 inhibitor NP3-253 (Supplementary Figures 4D and 4E). These results demonstrate that the incomplete inhibition of IL-1β following treatment with NK7-902 in human whole blood is not due to reduced activity of the compound in neutrophils.

The role of NEK7 in cell cycle progression raises a significant risk when considering potential NEK7- directed therapeutics. As defects in cell cycle progression and mitosis could eventually lead to chromosomal damage, we performed a micronucleus test in human primary blood lymphocytes treated with NK7-902 and found no significant increase in the number of micronucleated binucleates (MNCB) versus DMSO-treated cells, despite a complete loss of the NEK7 protein (Supplementary Figure 5A and 5B). Consistent with this, no obvious cell cycle progression defects were observed in *Nek7* KO HeLa and U2-OS cells compared to WT cells (Supplementary Figure 6).

### NK7-902 degrades NEK7 in mouse cells and in mice

IMiDs and other CRBN glue degraders are reportedly inactive in mouse cells, due to a single amino acid substitution, where the larger isoleucine 391 of mouse CRBN interferes with interaction occurring between valine 388 of human CRBN and the neosubstrates ^36^. Surprisingly, when we tested NK7-902 in mouse primary splenocytes, we observed that NK7-902 induces profound degradation of endogenous mouse NEK7, although with reduced potency (∼30x higher DC_50_) relative to human primary cells (Figure 4A). NK7-902-mediated NEK7 degradation was CRBN-dependent as it was strongly reduced in the presence of competing concentrations of lenalidomide and blocked by the addition of the NEDD8 inhibitor MLN4924 (Figure 4B). Furthermore, quantitative proteomic analysis of mouse primary splenocytes treated with 1 µM NK7-902 for 18h revealed that NEK7 was the only significantly down-regulated protein in these cells (Supplementary Figure 7).

**Figure 4.**
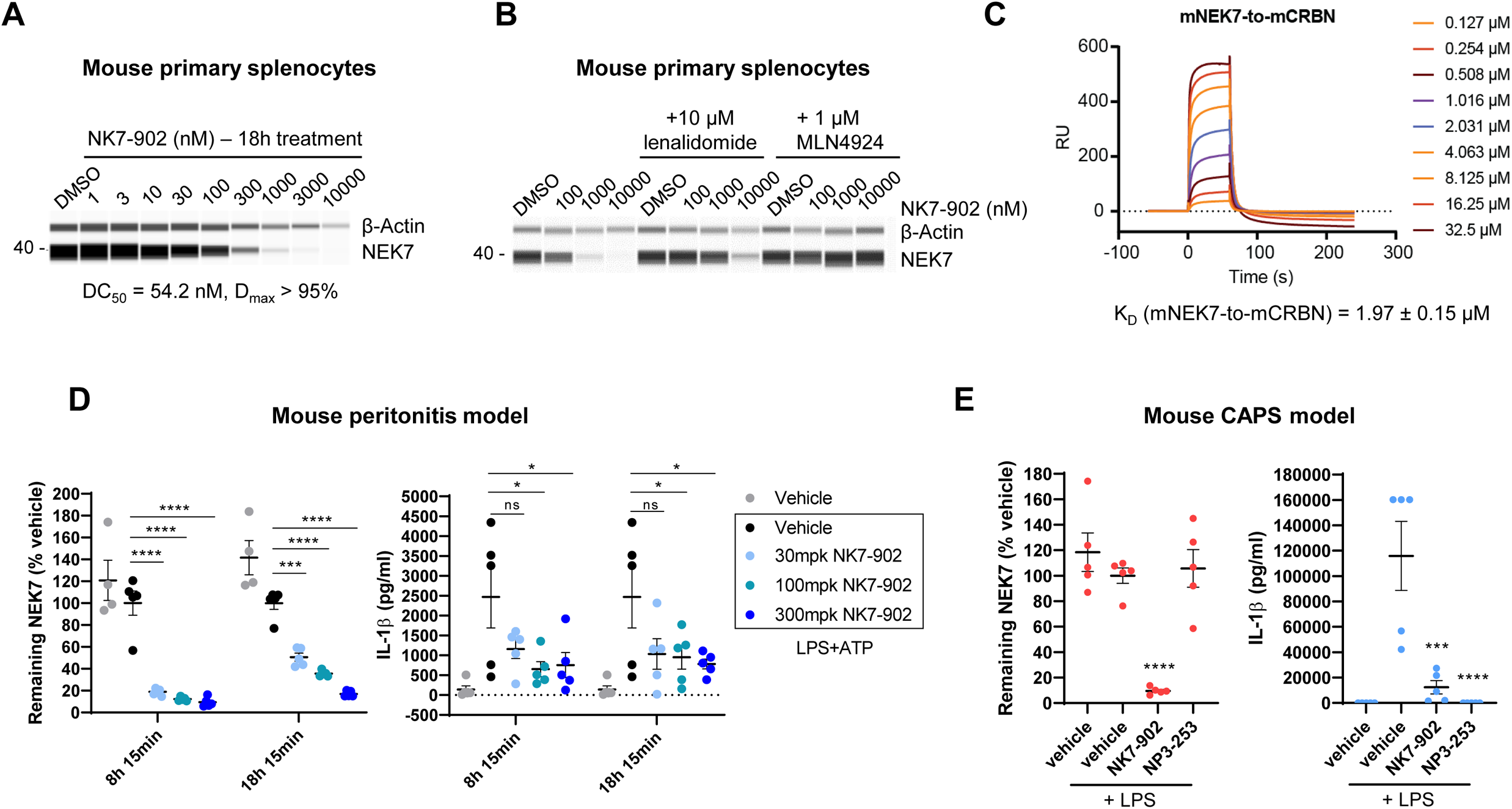
NK7-902 is active in mouse. (A) Mouse primary splenocytes were treated with increasing concentrations of NK7-902 for 18 h and then lysed to measure NEK7 and β-actin levels by WES capillary electrophoresis immunoanalysis. A representative experiment out of two independent repeats is shown. (B) Mouse primary splenocytes were incubated with the indicated concentrations of NK7-902 in the absence or presence of 10 µM lenalidomide (LFZ106) or the neddylation inhibitor MLN4924 for 18h. Cells were then lysed and NEK7 and β-actin levels were measured by WES. A representative experiment out of two independent repeats is shown. (C) SPR sensorgrams of the ternary complex of mNEK7 recruited to immobilized mCRBN//mDDB1 mediated by NK7-902. Shown are different concentrations of mNEK7 injected with a constant concentration of 5 µM NK7-902. The average KD and standard deviation of 2 independent experiments are shown below. (D) Mice (n=5 per group) were treated orally with either vehicle or increasing doses of NK7-902 for 6h or 16h and then challenged via intraperitoneal (i.p.) injection of LPS (2.5 mg/kg) for 2h followed by ATP (20 mM/0.5 mL/mouse) for an additional 15 min. NEK7 levels were measured in spleen lysates by WES, normalized over the corresponding β-actin signal and plotted (left). IL-1β levels were measured in the peritoneal lavage using a Bio-Rad multiplex immunoassay. (E) NLRP3 (A350V)/Cre+ mice (n=5 per group) were treated with 5 mg/kg tamoxifen, given i.p., for 5 days to induce expression of mutant NLRP3. Mice were then treated orally with vehicle or either 300 mg/kg NK7-902 or 20 mg/kg NP3-253 for 16h prior to being challenged with 10 µg/kg LPS i.p. for an additional 4h. A PBS-treated vehicle control group was also included to provide basal plasma IL-1β levels, which were measured using a Bio-Rad multiplex immunoassay. NEK7 levels were measured in spleen lysates by WES, normalized over the corresponding β-actin signal and plotted (left). Data in (D) and (E) represent mean ± SEM. Statistical significance was calculated using a one-way ANOVA with Dunnett’s multiple comparison test. **** p < 0.0001, *** p = 0.0001, * p < 0.05, ns = not significant.

Analysis of the hCRBN-NK7-902-hNEK7 ternary complex structure indicated that valine 388 is more solvent exposed at the CRBN-NEK7 protein-protein interface compared to the CRBN-lenalidomide-CK1α and CRBN-pomalidomide-IKZF1 ternary complex structures. Computationally introducing the V388I point mutation into the CRBN-NK7-902-NEK7 structure indicated that isoleucine may be accommodated without a steric clash, thus suggesting that recruitment can take place in mice (Supplementary Figure 8). We additionally characterized CRBN:NK7-902 interactions and CRBN:NK7-902:NEK7 ternary complexes using SPR. NK7-902 bound tightly to human DDB1:CRBN (K_D_ 49 nM) and also bound mouse DDB1:CRBN, but with weaker affinity (K_D_ 702 nM) (Supplementary Figure 9A). Using immobilized human DDB1:CRBN, excess (50 μM) NK7-902 was able to recruit human NEK7 with a K_D_ of 1.5μM (Supplementary Figure 9B). Meanwhile, using mouse DDB1:CRBN, excess NK7-902 recruited mouse NEK7 with a K_D_ of 2.6 μM, only marginally weaker than human (Figure 4C and Supplementary Figure 9B). Kinetic analysis also suggested a slightly shorter half-life of the mouse ternary complex compared to the human one (Supplementary figure 9B). These data support the ability of NK7-902 to function as a glue degrader of NEK7 in mouse cells. They further suggest that the reduced potency of NK7-902 induced NEK7 degradation in mice may result from decreased affinity of NK7-902 for mouse CRBN, as opposed to clashes between mouse CRBN and mouse NEK7.

NK7-902 had acceptable pharmacokinetic (PK) properties in mice with good oral exposure, enabling its use *in vivo* (Supplementary Figure 10). The activity of NK7-902 *in vivo* was first assessed in a mouse acute peritonitis model ^37^. Mice were pre-treated with single increasing oral doses of NK7-902 for either 6h or 16h and then challenged via intra-peritoneal injection of LPS for 2h, followed by a 15min ATP treatment to induce NLRP3 inflammasome activation. NK7-902 induced dose-dependent degradation of NEK7 in mouse spleen *in vivo*, with maximum effect observed with the 300 mg/kg dose at the 8h + 15min time point (Figure 4D). A slight recovery of NEK7 protein levels was seen at the later 18h + 15min time point. Importantly, the NK7-902-mediated NEK7 degradation was accompanied by a dose-dependent inhibition of IL-1β release upon *in vivo* activation of the NLRP3 inflammasome with LPS plus ATP (Figure 4D).

The activity of NK7-902 was also tested in the mouse NLRP3^A350V^ Muckle-Wells Syndrome (MWS) model of CAPS. Mice expressing the constitutively active NLRP3 ^A350V^ mutant, corresponding to the classical A352V MWS in human NLRP3, were first treated for 5 days with tamoxifen to induce expression of the NLRP3 mutant protein. Mice were next dosed orally with the NEK7 degrader (300 mg/kg NK7-902) or a NLRP3 inhibitor (20 mg/kg NP3-253) for 16h and then challenged with LPS for 4h to activate the inflammasome. NK7-902 strongly degraded NEK7 in the spleen of these mice and inhibited IL-1β release in blood to levels approaching the full inhibition seen with the NLRP3 inhibitor (Figure 4E).

These data demonstrate that NK7-902 is capable of degrading mouse NEK7 *in vivo*, thereby inhibiting the function of the NLRP3 inflammasome in an LPS/ATP peritonitis model and a CAPS disease mouse model.

### NEK7 degradation by NK7-902 *in vivo* in non-human primates (NHP) leads to incomplete and transient IL-1β inhibition

Due to the high doses of NK7-902 needed to induce NEK7 degradation in mouse and the known differences in the off-target profile of CRBN glue degraders in rodents versus primates, cynomolgus monkey (also referred to as NHP in our manuscript) was selected as a species for subsequent pre-clinical work.

The activity of NK7-902 was first assessed in cynomolgus monkey blood *in vitro.* NK7-902 potently degraded NEK7 and partially inhibited IL-1β release in NHP blood (Supplementary Figure 11A and 11B) similar to what was observed in human blood (Figure 3D and 3E). NK7-902 PK and pharmacodynamics (PD) was assessed in NHP *in vivo* following a single intravenous dose of 1 mg/kg or single oral doses of either 0.2 or 2 mg/kg. NK7-902 showed moderate clearance (26.1 mL/min/kg) following intravenous administration and an over-proportional exposure between 0.2 and 2 mg/kg oral dosing based on mean dose normalized C_max_ and AUC_last_ values. Oral bioavailability was 14.3% following oral gavage administration of 0.2 mg/kg and 49.7% with 2 mg/kg compared with the intravenous dose at 1 mg/kg (Supplementary Figure 12). Oral administration of NK7-902 *in vivo* led to dose-dependent and sustained NEK7 degradation in blood (Figure 5A and 5B). The 0.2 mg/kg dose induced a maximum of 70% NEK7 degradation, with a small recovery of NEK7 levels observed only 72h post dose (Figure 5A). More than 95% NEK7 degradation was seen with the 2 mg/kg dose 2h post-dose and these low NEK7 levels were maintained up to 48h post-dose (Figure 5B). Importantly, NEK7 degradation in blood persisted in the absence of detectable levels of NK7-902 at both doses, highlighting the expected disconnect between PK and PD previously reported with other degrader molecules ^38^.

**Figure 5.**
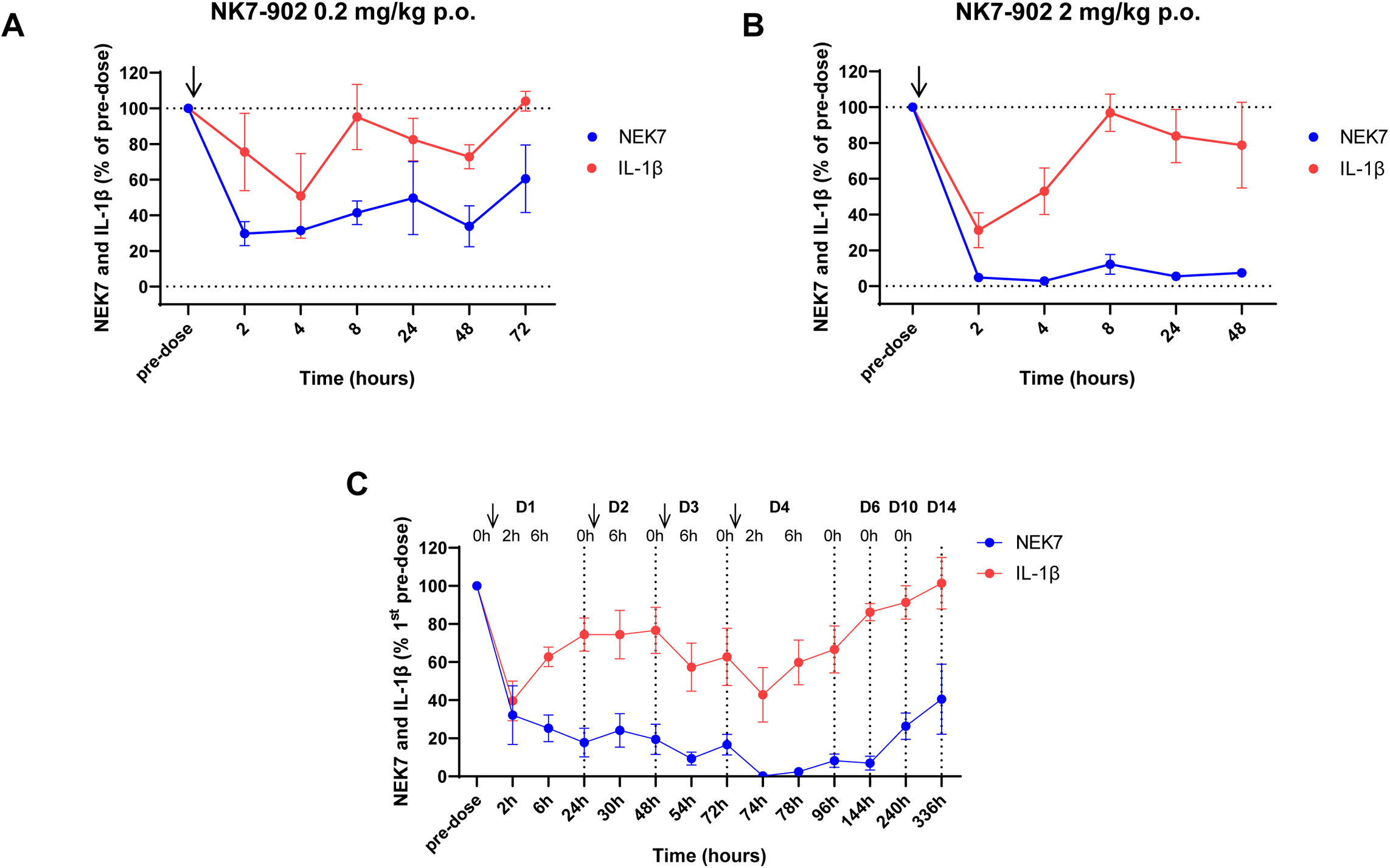
NK7-902 degrades NEK7 but only partially inhibits IL-1β release *in vivo* in non-human primates. (A) Three NHPs were given one dose of NK7-902 at either 0.2 mg/kg or 2 mg/kg orally and blood samples were collected at the indicated times. Red blood cells were lysed and NEK7 and β-actin levels were measured in whole blood cell protein extracts by WES capillary electrophoresis immunoanalysis. (B) Blood samples were collected from the same animals as in (A) and stimulated *ex-vivo* with 1 ng/mL LPS for 3h, followed by an additional 30min with 3 mM ATP. IL-1β levels in the plasma fraction were measured by HTRF. (C) Three NHP were given four consecutive daily oral doses of NK7- 902 at 0.2 mg/kg and two identical sets of blood samples were collected at the indicated times (pre-dose, 2h and 6h post dose for the first four days and then pre-dose on day 5, 6, 10 and 14). One set was processed for red blood cell lysis and then NEK7 and β-actin levels were measured in white blood cell protein extracts by WES capillary electrophoresis immunoanalysis. The second set of blood samples were stimulated *ex-vivo* with 1 ng/mL LPS for 3h, followed by an additional 30min with 3 mM ATP. IL-1β levels in the plasma fraction were measured by HTRF. Data in (A-C) represent mean ± SEM. Arrows indicate oral administration of NK7-902.

To study NLRP3 inflammasome inhibition following *in vivo* NEK7 degradation in NHP, blood was stimulated *ex vivo* with LPS and ATP to induce IL-1β release. The *ex vivo* stimulation protocol was first tested in a mouse experiment and proved to work successfully, with NEK7 degradation *in vivo* leading to sustained inhibition of IL-1β in *ex vivo* stimulated mouse blood (Supplementary Figure 11C). A maximum of 70% IL-1β inhibition was measured 2h after oral administration of NK7-902 at 2 mg/kg in NHP (Figure 5B). IL-1β levels increased again rapidly at later time points, reaching values similar to those measured in the pre-dose samples from 8h onwards (Figure 5B).

To confirm and expand on these findings, we performed a second NHP study where NK7-902 was dosed orally for four consecutive days at 0.2 mg/kg (Figure 5C and Supplementary Figure 12B). Stepwise and progressive NEK7 degradation was detected after multiple doses of NK7-902, leading to no detectable NEK7 protein 2h after the last dose on day 4. Partial recovery of NEK7 levels was observed starting from 168h after the fourth dose (day 10), suggesting a long NEK7 protein half-life *in vivo*. Despite profound and stable NEK7 degradation upon treatment with NK7-902, IL-1β release upon stimulation *ex vivo* was efficiently inhibited only very transiently (mostly visible at 2h after the first dose) and then rose again, consistent with the observations made in the single dose NHP experiment (Figure 5C).

Together, these data demonstrate that NK7-902 is a potent degrader of NEK7 in NHP *in vivo*, capable of sustained and efficient NEK7 degradation with once daily dosing. The IL-1β results, however, also highlight a disconnect between NEK7 degradation and inhibition of NLRP3-dependent IL-1β release. These results corroborate our findings in human whole blood and suggest alternative mechanisms for NLRP3 pathway activation in the absence of NEK7 in non-human primates.

## Discussion

In this report, we describe the discovery of NK7-902, a potent, selective, and orally bioavailable CRBN glue degrader of NEK7, and characterize its potential to treat NLRP3-driven diseases. A glue degrader modality represents an attractive approach to target NEK7 in the context of the NLRP3 inflammasome because NLRP3 requires the scaffolding function of NEK7, which may not be blocked by a NEK7 kinase inhibitor. We found that NK7-902 degrades NEK7 highly efficient *in vitro* and demonstrated, through structural work, point mutation studies and CRBN knockout experiments, that this requires both the NEK7 β-hairpin and CRBN. We showed that the extent to which NEK7 degradation affected NLRP3-mediated IL-1β release was variable, depending on the donor and the experimental conditions. In human primary monocytes, NEK7 depletion inhibited IL-1β release by 79% and 75% upon stimulation with LPS plus ATP or with niclosamide, respectively, but it only minimally affected the release of this cytokine upon nigericin activation. Similarly, in human whole blood stimulated with LPS/ATP, NK7-902-mediated depletion of NEK7 caused only 60% inhibition of IL-1β release. To further probe the reduced effect on IL-1β release by NEK7 degradation in whole blood compared to monocytes, we isolated neutrophils, which are important producers of IL-1β besides monocytes ^39,40^. However, testing the effect of NEK7 degradation showed that NK7-902 can effectively block the NLRP3 pathway in isolated neutrophils. Collectively, the data suggested that NEK7 is not always required for NLRP3 activation and that additional factors may substitute for NEK7 or compensate in the absence of NEK7.

Initial structural work examining NEK7 in complex with inactive NLRP3 suggested a model where NEK7 provides a ‘license’ for NLRP3 activation, enabling its oligomerization into an active inflammasome complex ^15^. In this model, NEK7 first interacts with the LRR and NACHT domains of inactive NLRP3, a process that may occur at the MTOC where NEK7 localizes and where NLRP3 is transported from the TGN38+ in the form of inactive cages for activation ^21^. Binding to NEK7, along with conformational changes of the NLRP3 NACHT domain, subsequently promote the formation of an active NLRP3 oligomer, where NEK7 is required to connect adjacent NLRP3 molecules by mediating interactions between NLRP3 LRR domains ^15^.

This model, which positions NEK7 as an indispensable factor for NLRP3 oligomerization, was challenged by the finding that, in the active NLRP3 inflammasome disc, the C-terminal LRR domain of NLRP3 as well as LRR-bound NEK7 extend away from the center of the disc and do not participate in disc interfaces ^24^. This finding, together with data demonstrating that a NLRP3 mutant lacking the LRR domain can still be activated ^26^, suggests that the interaction with NEK7 is not strictly needed for NLRP3 inflammasome oligomerization. Xiao et al. proposed that NEK7 may instead serve to break the inactive NLRP3 cages at the MTOC, promoting opening of the cages and their rearrangement into active NLRP3 oligomers. However, in conditions associated with high NLRP3 expression or destabilization of the inactive NLRP3 cage, inflammasome assembly may occur from either monomeric or caged NLRP3 independently of NEK7 ^24^.

In line with these findings, the NLRP3 inflammasome was shown to be fully functional in *Nek7* KO iPS-derived human macrophages which, unlike murine macrophages, appear to rely mostly on a IKKβ- dependent pathway for NLRP3 activation ^22^. Mechanistically, IKKβ primes NLRP3 by increasing its recruitment to phosphatidylinositol 4-Phosphate (PI4P)-containing membranes. Some dependency on NEK7 was observed following short stimulations *in vitro* but with longer stimulations NEK7 appeared to be fully dispensable for NLRP3 activation. It was proposed that NEK7 may accelerate inflammasome formation initially but becomes redundant once the IKKβ priming cascade is fully operational ^22^. In fact, there is evidence that pharmacological inhibition of IKKβ can block NLRP3 activation in THP-1 cells independently of transcriptional priming ^41,42^, which remains challenging to test in human primary monocytes, PBMCs and whole blood where an IKKβ-dependent transcriptional step is essential for IL-1β production. Thus, it remains possible that an IKKβ-dependent but NEK7-independent pathway contributes to IL-1β secretion in our human and monkey cytokine experiments.

A context-dependent role for NEK7 is further supported by the recent discovery that NLRP3 can be activated through two parallel and biologically distinct pathways in human THP1 cells: the previously described MTOC-dependent pathway, requiring NLRP3 oligomerization into barrel/cage-like structures; and a second MTOC-independent one, involving monomeric NLRP3 species that do not form cages ^25^. Both pathways appear to be functional upon stimulation by, *e.g.*, nigericin, although the MTOC-associated NLRP3 form executes inflammasome functions, such as IL-1β release and pyroptosis, more quickly ^25^. Stimuli such as imiquimod, by contrast, only engage the MTOC associated pathway. Remarkably, deficiency in NEK7 was previously shown to abolish imiquimod-dependent NLRP3 activation, whereas inhibition of nigericin- or ATP-driven NLRP3 activation was only partial under those conditions ^22^. This is consistent with our data and with the hypothesis that NEK7 is required for the MTOC-dependent NLRP3 activation pathway, but dispensable in the context of MTOC-independency. Kinetics data showing that NEK7 accelerates NLRP3 activation ^22^ agree with a selective role of NEK7 in the MTOC-dependent pathway, which has an early onset compared to the MTOC-independent one ^25^. Of note, beyond NEK7, other NLRP3 regulators are emerging to be selectively involved in the MTOC-dependent pathway. For instance, protein kinase D, a key kinase for licensing of the NLRP3 inflammasome ^42,43^ was shown to accelerate NLRP3 activation by nigericin and to be dispensable at later time point ^44^. The recently reported context-dependent requirement for HDAC6 also speaks for its selective involvement in the MTOC-dependent NLRP3 activation pathway ^24,25,45,46^.

Unlike other reported CRBN-based glue degraders, NK7-902 was able to degrade NEK7 in mouse cells (albeit with reduced potency), allowing us to explore the effects of NEK7 degradation in mice. We found that NK7-902 mediated NEK7 degradation leads to sustained inhibition of NLRP3 dependent IL-1β release in the peritonitis and CAPS mouse models, consistent with data obtained in macrophages from Nek7 KO mice, which supports a key role for NEK7 in NLRP3 activation, as reported by several groups ^12–14^. Several lines of evidence suggest that NLRP3 activation in murine cells may be selectively wired towards MTOC-dependency. Microtubule disruption, for instance, could abolish nigericin induced NLRP3 activation in immortalized mouse bone marrow-derived macrophages ^45^ whereas it only partially impacted human NLRP3 activation in THP-1 cells ^25^. In addition, murine NLRP3 appears to be more promiscuous than human NLRP3 in terms of lipid binding capacity; it can interact with phosphatidic acid (PA) as well as all phosphorylated phosphatidylinositols (PtdIs), whereas human NLRP3 only binds PA and PtdIns (3,4,5)-P3 ^25^. This further supports the more stringent MTOC pathway dependency of murine NLRP3, for which interaction with membranes is critical.

In our PK/PD experiments in non-human primates, we found that blockade of IL-1β release after NLRP3 activation was transient and partial despite profound and long-lasting down-regulation of NEK7 protein levels. This is in line with our findings in human and cynomolgous blood *in vitro* and further suggests a degree of plasticity in NLRP3 activation mechanisms whereby depletion of NEK7 may shift inflammasome activation towards alternative non-NEK7 dependent mechanisms. Several factors, such as the species, the stimulus, the cell type and its physiological state, the available pool of NLRP3 at the membrane (caged) vs. cytosolic NLRP3 (non-oligomeric), and the NLRP3 interactome, including post-translational modifiers and their activation status, likely all have an impact to modulate the path for NLRP3 activation^42^.

In our hands, NEK7 knockout or degradation did not lead to obvious cell cycle or chromosomal defects. This is in contrast with what was previously described, even though we specifically selected HeLa and U2OS cells for knocking-out NEK7 as most of the studies addressing the role of NEK7 in cell cycle were performed using these two cell lines ^9,11,47,48^. It should be noted that the cell cycle defects observed in these reports are generally very subtle, with only a small proportion of cells showing the phenotype. Importantly, mouse-derived embryonic fibroblasts from Nek7 KO mice developed a binuclear phenotype that was significantly more pronounced compared to control cells only after multiple passages *in vitro* ^49^. This suggests that longer cultures may be needed for detecting differences between WT and Nek7 KO or NK7-902 treated cells. Additional studies should be conducted to further evaluate potential cell-cycle associated safety risks following NEK7 degradation.

Overall, our data demonstrate a context- and species-specific role for NEK7 in the NLRP3 inflammasome, confirming previous observations and further contributing to our understanding of the mechanisms regulating NLRP3 activation in human cells. While these results question the applicability of a NEK7 degrader molecule for the treatment of NLRP3-driven diseases, NK7-902 represents a unique chemical tool to continue explore the function of NEK7 in the NLRP3 pathway and beyond.

### Significance

Hyperactivation of the NLRP3 inflammasome has been linked to a variety of acquired and inherited diseases, which has prompted the search for therapeutics that can inhibit the NLRP3 pathway. While direct NLRP3 small molecule inhibitors are currently being tested in the clinic, the mitotic kinase NEK7 has recently emerged as a potential alternative target to block NLRP3 activity. As the function of NEK7 in the NLRP3 inflammasome is independent of its kinase activity, we hypothesized that degradation of NEK7 with a glue degrader molecule could be a unique approach to inhibit the pathway. We identified NK7-902, a CRBN glue degrader of NEK7, capable of inducing profound degradation of NEK7 in human and monkey primary cells and blood. Unlike other CRBN-based glue degraders, NK7-902 was also able to effectively degrade NEK7 in mice, albeit with reduced potency. Unexpectedly, treatment with NK7-902 only partially and transiently blocked NLRP3-dependent IL-1β release in human and cynomolgus cells/blood, as opposed to the full inhibition seen with a direct NLRP3 inhibitor. Our results indicate that, in these species, the NLRP3 inflammasome can, at least in part, be activated in a NEK7-independent manner, questioning the applicability of a NEK7 degrader molecule for the treatment of NLRP3-driven diseases. At the same time, our work contributes significantly to understanding the role of NEK7 in different species and conditions, and reveals the structure of a unique, orally bioavailable, chemical tool to continue explore the function of NEK7 in the NLRP3 pathway and beyond.

## Acknowledgements

The authors thank many current or past Novartis colleagues whose efforts supported this work: Carina Sanchez and Shan Yang for their help in material supply, Nicole von Burg and Joerg Eder for careful reading and editing of the manuscript, Christina Hebach for helpful advice during the execution of the project, Gregory Hollingworth, William Forrester, Rohan Beckwith, Rishi Jain, Richard Siegel, Christian Bruns, James Bradner, John Tallarico for their intellectual input and strategic support. A. N. W. R. gratefully acknowledges infrastructural support provided by the University of Tübingen, the University Hospital Tübingen and the DFG Clusters of Excellence “iFIT – Image-Guided and Functionally Instructed Tumor Therapies” (EXC 2180) and “CMFI – Controlling Microbes to Fight Infection (EXC 2124). This work was funded by Novartis Biomedical Research.

## Author contributions

F.M., A.C. and D.L.B. designed, synthesized and/or characterized the reported compounds, A.B., P.R., J.S., J.P., L.X. and A.S. developed and performed biochemical and cellular assays to support compound characterization, studies in primary cells and ex-vivo work was performed by A.S., N.S and T.B. Y.C.G, S.B., A.N.R.W., T.W. and J.K.D collected and provided PBMCs from CAPS patients. M.M., D.B., P.L., E.A. and A.H. conducted expression proteomics experiments, M.S., M.K., J.L. and C.W. supported and performed protein biophysics and cryo-EM structural work. C.J.D. aided in modeling NK7-902 with CRBN and NEK7. M.F., A.E., M.B. performed the micronucleus test in primary lymphocytes, S.M and S.L. performed cell cycle studies in Nek7 KO cells. M.V., M.S., J.D. designed and performed in vivo mouse experiments, A.Z., M.K. and P.A. were responsible for pharmacokinetic studies in mouse and NHP, D.L.B. and Z.I.B. led the project, M.S., J.S., F.B., D.L.B. and Z.I.B. wrote and edited the manuscript, J.S., C.F. and R.P. contributed with intellectual and strategic input.

## Inclusion and diversity

We support inclusive, diverse and equitable conduct of research.

## Supplemental information

Supplemental information includes the following:

Supplementary Figures. Figures S1 – S12

Supplementary Tables. Tables S1 – S3 containing complete proteomics data related to Figure 2C, S3B and S7.

## Resource availability

### Lead contact

Further information and requests for resources and reagents should be directed to the lead contact, Zuni Bassi.

### Materials availability

All unique/stable reagents generated in this study are available from the lead contact with a completed materials transfer agreement.

### Data and code availability

The refined EM map and the coordinates of the CRBN-NK7-902-NEK7 complex have been deposited in the PDB (pdb ID:9H59) and EMDataBank (ID: EMD-51881). The mass spectrometry proteomics data have been deposited to the ProteomeXchange Consortium via the PRIDE partner repository and reviewer access will be provided upon request. This paper does not report original code.

## Materials and Methods

### NEK7 Compounds

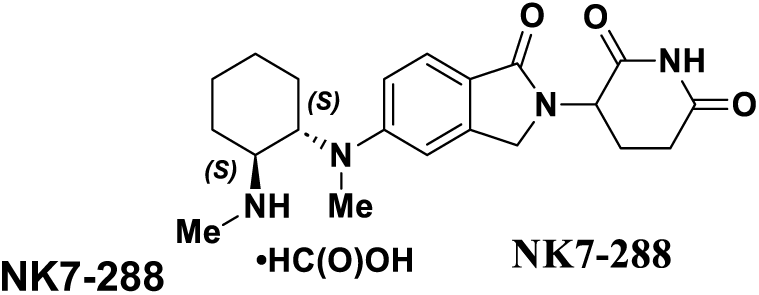

NK7-288 was synthesized as described in US2020/16143, 2020.

#### 2-(2,6-dioxopiperidin-3-yl)-5-(methyl((1S,2S)-2-(methylamino)cyclohexyl)amino)isoindoline-1,3- dione (NK7-147)

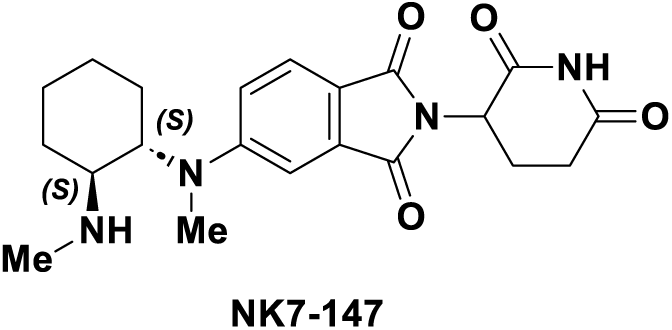

**NK7-147** was synthesized as described in US2020/16143, 2020.

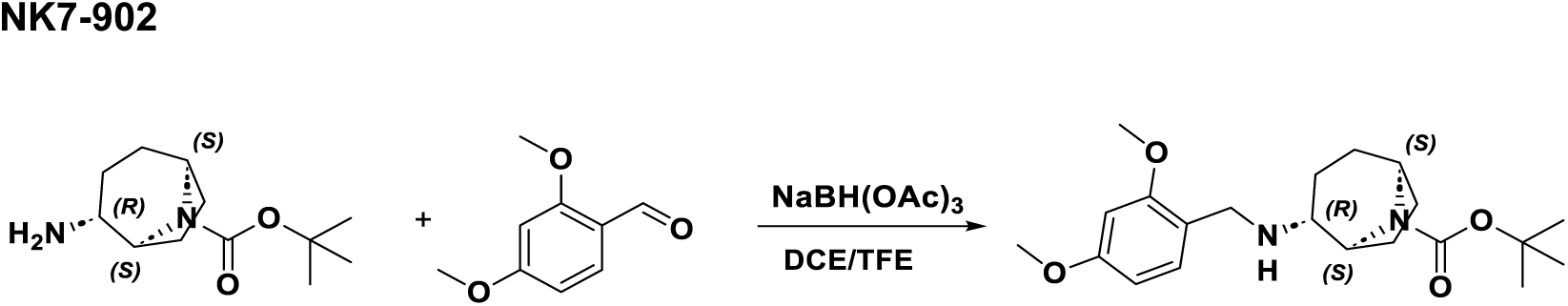

#### *tert-*butyl (1S,2R,5S)-2-((2,4-dimethoxybenzyl)amino)-8-azabicyclo[3.2.1]octane-8-carboxylate

Acetic acid (0.076 mL, 1.3 mmol) was added to the mixture of *tert-*butyl (1S,2R,5S)-2-amino-8- azabicyclo[3.2.1]octane-8-carboxylate (3.00 g, 13.3 mmol) and 2,4-dimethoxybenzaldehyde (11.01 g, 66.3 mmol) in DCE (70 mL) and TFE (70 mL), followed by sodium triacetoxyborohydride (19.67 g, 93 mmol). The resulting mixture was stirred overnight and then concentrated. The residue was partitioned between DCM and saturated sodium bicarbonate solution and the aqueous phase was washed with DCM (3x). The combined organic phases were dried over anhydrous sodium sulfate, filtered, and concentrated. The residue was purified via flash chromatography (eluting with0-100% EtOAc/ Heptane) to provide desired product (4.8 g, 12.8 mmol, 96%) as a white solid. ^1^H NMR (400 MHz, MeOD) δ 7.37 (d, *J* = 8.4 Hz, 1H), 6.66 (d, *J* = 2.4 Hz, 1H), 6.61 (dd, *J* = 8.3, 2.4 Hz, 1H), 4.76 – 4.68 (m, 1H), 4.46 (dt, *J* = 6.2, 2.9 Hz, 1H), 4.29 (q, *J* = 13.1 Hz, 2H), 3.93 (s, 3H), 3.85 (s, 3H), 3.23 (t, *J* = 3.9 Hz, 1H), 2.17 – 1.72 (m, 8H), 1.63 – 1.55 (m, 1H), 1.53 (s, 9H). MS [M+H]^+^ = 377.3.

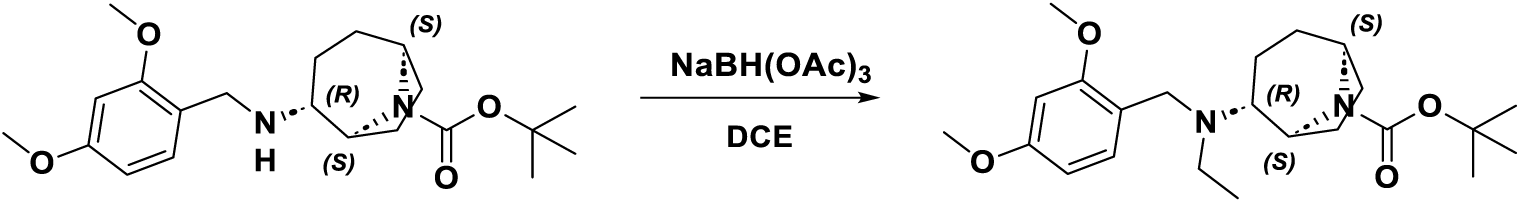

#### *tert-*butyl (1S,2R,5S)-2-((2,4-dimethoxybenzyl)(ethyl)amino)-8-azabicyclo[3.2.1]octane-8- carboxylate

Acetic acid (0.073 mL, 1.3 mmol) was added to a mixture of *tert-*butyl (1S,2R,5S)-2-((2,4- dimethoxybenzyl)amino)-8-azabicyclo[3.2.1]octane-8-carboxylate (4.8 g, 13 mmol) and acetaldehyde (3.58 mL, 63.7 mmol)) in DCE (150 mL), followed by sodium triacetoxyborohydride (16.21 g, 76 mmol). The resulting mixture was stirred overnight and then concentrated. The residue was partitioned between DCM and saturated sodium bicarbonate solution and the aqueous phase was washed with DCM (3x). The combined organic phases were dried over anhydrous sodium sulfate, filtered, and concentrated to provide crude desired product (4.80 g, 11.9 mmol, 93%) as a white solid. ^1^H NMR (400 MHz, MeOD) δ 7.33 (d, *J* = 8.1 Hz, 1H), 6.48 (d, *J* = 8.3 Hz, 2H), 4.58 (d, *J* = 7.1 Hz, 1H), 4.32 (d, *J* = 6.3 Hz, 1H), 3.79 (s, 3H), 3.78 (s, 3H), 3.77 (s, 2H), 2.86 – 2.73 (m, 2H), 2.70 – 2.62 (m, 1H), 1.99 – 1.80 (m, 4H), 1.77 – 1.60 (m, 3H), 1.47 (s, 9H), 1.34 (t, *J* = 10.5 Hz, 1H), 1.01 (t, *J* = 7.1 Hz, 3H). MS [M+H]^+^ = 405.2.

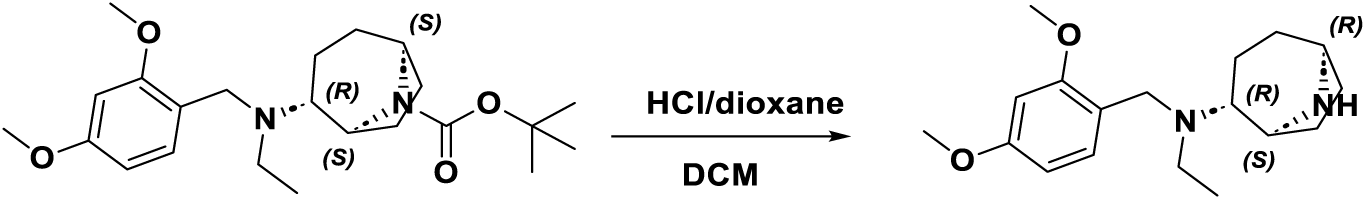

#### (1S,2R,5R)-N-(2,4-dimethoxybenzyl)-N-ethyl-8-azabicyclo[3.2.1]octan-2-amine

4M HCl in dioxane (14.52 mL, 58.1 mmol) was added to a mixture of *tert-*butyl (1S,2R,5S)-2-((2,4- dimethoxybenzyl)(ethyl)amino)-8-azabicyclo[3.2.1]octane-8-carboxylate (4.7 g, 11.62 mmol) in DCM (150 mL). The resulting mixture was stirred overnight and then concentrated to dryness to provide crude desired product as the HCl salt (4.7 g, 11.2 mmol, 96%) as a white solid. MS [M+H]^+^ = 305.2.

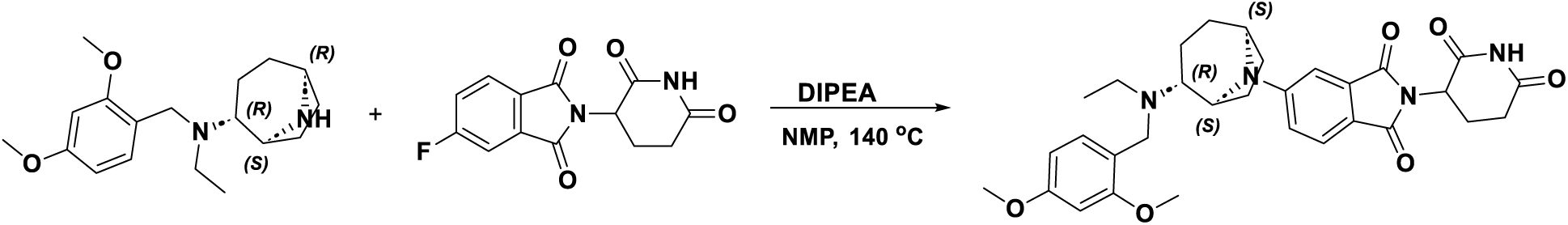

#### 5-((1S,2R,5S)-2-((2,4-dimethoxybenzyl)(ethyl)amino)-8-azabicyclo[3.2.1]octan-8-yl)-2-(2,6- dioxopiperidin-3-yl)isoindoline-1,3-dione

A mixture of 2-(2,6-dioxopiperidin-3-yl)-5-fluoroisoindoline-1,3-dione (4.28 g, 15.50 mmol), (1S,2R,5R)-N- (2,4-dimethoxybenzyl)-N-ethyl-8-azabicyclo[3.2.1]octan-2-amine hydrogen chloride salt (4.50 g, 11.93 mmol), and DIPEA (20.83 mL, 119 mmol) in NMP (20 mL) was stirred at 140 °C o/n. The resulting mixture was diluted with water and the aqueous phase was washed with DCM.The organic phase was washed with water and brine, dried over anhydrous sodium sulfate, filtered, and concentrated. The crude residue was purified via reverse phase C18 chromatography, eluting with 5-100% 0.1% TFA in ACN/water. The desired fractions were collected, neutralized with saturated sodium bicarbonate solution, the product was extracted with DCM, and then further purified via flash chromatography (eluting with 0-30% 2% TEA in mixture of EtOAc and EtOH (4:6)/ DCM) to provide desired product (5.8 g, 10.35 mmol, 87%) as a light yellow powder. ^1^H NMR (400 MHz, MeOD) δ 7.53 (dd, *J* = 8.5, 4.0 Hz, 1H), 7.10 (dd, *J* = 7.9, 2.3 Hz, 1H), 6.95 (dd, *J* = 8.5, 2.3 Hz, 1H), 6.77 (dd, *J* = 13.0, 8.4 Hz, 1H), 6.37 (t, *J* = 2.5 Hz, 1H), 6.24 (ddd, *J* = 8.4, 4.4, 2.4 Hz, 1H), 5.03 (dd, *J* = 12.4, 5.5 Hz, 1H), 4.68 (d, *J* = 6.3 Hz, 1H), 4.35 (s, 1H), 3.71 (d, *J* = 3.0 Hz, 6H), 3.67 – 3.48 (m, 2H), 2.86 (ddd, *J* = 17.6, 14.1, 5.1 Hz, 1H), 2.78 – 2.52 (m, 5H), 2.15 – 2.05 (m, 1H), 1.97 (dq, *J* = 15.7, 4.7 Hz, 3H), 1.91 – 1.72 (m, 4H), 1.39 (dt, *J* = 11.2, 5.7 Hz, 1H), 0.92 (t, *J* = 6.9 Hz, 3H). MS [M+H]^+^ = 561.6.

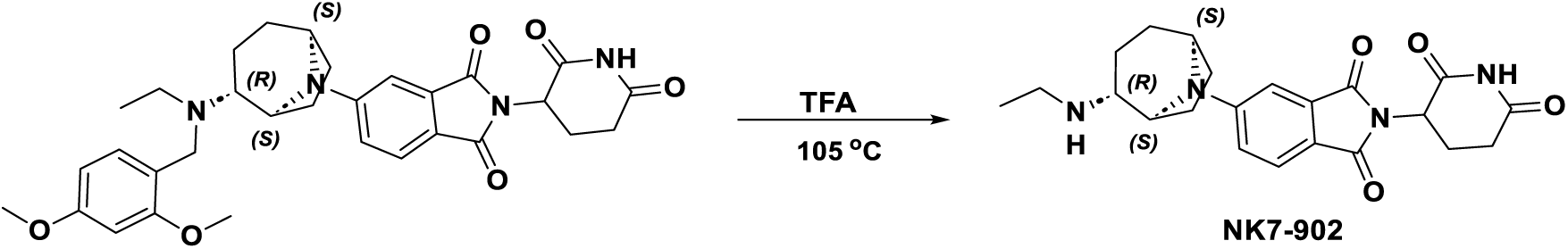

#### 2-(2,6-dioxopiperidin-3-yl)-5-((1S,2R,5S)-2-(ethylamino)-8-azabicyclo[3.2.1]octan-8-yl)isoindoline-1,3-dione (NK7-902)

A mixture of 5-((1S,2R,5S)-2-((2,4-dimethoxybenzyl)(ethyl)amino)-8-azabicyclo[3.2.1]octan-8-yl)-2-(2,6- dioxopiperidin-3-yl)isoindoline-1,3-dione (5.8 g, 10.35 mmol) in TFA(100 mL) was stirred at 105 °C for 3 hrs. The reaction mixture was concentrated and the resulting residue was partitioned between DCM and saturated sodium bicarbonate solution. The product was extracted with DCM (3x) and the combined organic phases were dried over anhydrous sodium sulfate, filtered, and concentrated. The residue was purified via reverse phase C18 chromatography, eluting with 5-100% 0.1% TFA in ACN/water. The desired fractions were collected, neutralized with saturated sodium bicarbonate solution, the product was extracted with DCM, and then further purified via flash chromatography (eluting with 0-30% 2% TEA in mixture of EtOAc and EtOH (4:6)/ DCM) to provide desired product **1** (3.3 g, 7.88 mmol, 76%) as a light yellow powder. MS [M+H]^+^ = 411.2. ^1^H NMR (acetonitrile-*d_3_*, 400 MHz) δ 8.6-9.3 (br s, 1H), 7.56 (d, *J = J =* 8.3 Hz, 1H), 7.15 (d, *J =* 2.4 Hz, 1H), 7.00 (dd, *J =* 2.2, 8.6 Hz, 1H), 4.9-5.1 (m, 1H), 4.55 (br d, *J =* 4.9 Hz, 1H), 4.3-4.4 (m, 1H), 2.5-2.9 (m, 7H), 2.1-2.2 (m, 4H), 1.8-1.9 (m, 2H), 1.4-1.6 (m, 2H), 0.91 (t, *J =* 7.1 Hz, 3H).

#### NP3-253 (R)-2-(6-((1-ethylpiperidin-3-yl)amino)pyridazin-3-yl)-3-methyl-5-(trifluoromethyl)phenol (NP3-253)

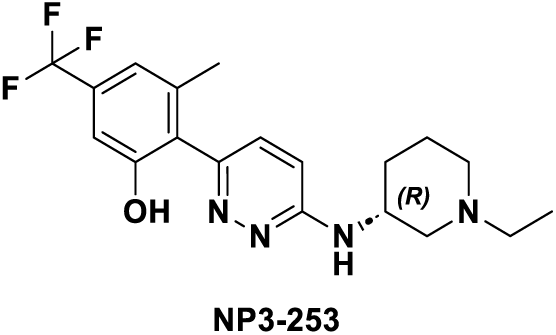

NP3-253 was synthesized as described in US2020/361898, 2020.

### TR-FRET biochemical recruitment assay

Forty nL of increasing concentrations of compounds diluted in dimethyl sulfoxide (DMSO) were dispensed directly into the wells of a white low volume 384 well plate via acoustic transfer (Beckman, ECHO 550R series) and incubated with 10 µL of a protein mix, dispensed with Certus (Fritz Gyger Ag, CERTUS nano liquid dispenser) and containing the following: 50 nM eGFP-labeled humanCRBN/humanDDB1 (produced in house), 50 nM de-phosphorylated/biotinylated NEK7 (produced in house) and 1 nM Terbium-Streptavidin (Invitrogen, PV3576) in buffer containing 50 mM TRIS-HCL, pH7.4, 100 mM NaCl, 2 mM EDTA, 2% Glycerol, 0.1% Pluronic F-127 and 1 mM TCEP. Following a 2min incubation, FRET emission of eGFP fluorescence was recorded using a Pherastar FSX instrument.

### Virus production and transduction

Lentivirus encoding Prolabel-NEK7 was made as follows: 4×10^6^ low-passage HEK293T cells were plated in 10 cm^2^ tissue culture plates in 10 mL complete Dulbecco’s modified Eagle’s medium (DMEM, ThermoFisher 10564011) growth media (10% FBS, 1% penicillin-streptomycin) and incubated overnight (37°C, 5% CO_2_). The following day, culture media was removed by aspiration and replaced with 6 mL DMEM. For each plate, 36 µL FugeneHD (Promega E2311) was added to 600 µL Opti-MEM™ (ThermoFisher 31985062), while 9 µg of ViraPower Packaging Mix (ThermoFisher K497500) and 3 µg of pLenti6.2-V5-Prolabel-NEK7 DNA was added to 50 µL Opti-MEM. After incubation at room temperature (RT) for 5 minutes, diluted DNA and Fugene mixtures were combined, incubated for 30 minutes at RT, and then added dropwise to cells prior to overnight incubation (37°C, 5% CO_2_). Transfection media was replaced with 6 mL fresh growth media the following day. Seventy-two hours post-transfection, viral supernatant was harvested, cleared by centrifugation at 418 x *g* for 10 minutes, and filtered through a 0.45 µM membrane syringe filter. Viral titer was approximated using Lenti-X GoStix (Takara 631280) and stored at 4°C prior to infection.

To transduce HEK293T with Prolabel-NEK7 virus, 500 µL of viral supernatant was added to 3×10^6^ cells in a final volume of 3 mL growth media supplemented with 8 µg/mL polybrene (Millipore Sigma TR-1003-G) in a 12-well dish. Cells were spinfected by centrifugation at 742 x *g* for 1h at 22°C. Cells were returned to incubator for 24h prior to trypsinization and replating in a 10 cm^2^ dish with 10 mL growth media supplemented with 10 µg/mL blasticidin (ThermoFisher A1113903). After 5-7 days, surviving cells were trypsinized and single clones were isolated via limiting dilution in 96 well plates. Prolabel-NEK7 expression in clonal populations was confirmed by of DiscoverX PathHunter Prolabel Detection Reagent (User Manual 93-0180) in 384-well plates according to manufacturer’s protocol.

### NanoBiT recruitment assay

CRBN was cloned into Promega LgBiT Flexi vector pFC34K and NEK7 and IKZF2 were cloned into Promega SmBiT Flexi vector pFN35K following the Promega protocol (N2015, NanoBiT Protein:Protein Interaction System Technical Manual). All DNA inserts were sequence verified. 293T cells were transfected as follows: 20 µg of a mixture containing a 1 to 3 ratio of LgBiT CRBN and SmBiT NEK7/IKZF2 plasmid was added to 1 mL of OptiMEM medium (ThermoFisher 31985062), mixed with 60 µL FuGene HD transfection reagent (Promega, E2311) and incubated at RT for 20 min. 293T cells were trypsinized and diluted in growth medium to a concentration of 1.0×10^6^ cells/mL and the transfection reagents were added to 10 mL of cells. After mixing, a 20 mM stock (in DMSO) of NAE1 inhibitor MLN4924 (AdipoGen, AG-CR1-3703) was added to cells for a final concentration of 2 µM. Cells (20 µL per well) were plated into 384 well solid white plates (Corning, 3570) and incubated for 24h. Test compounds were serially diluted in DMSO and 100 nL of compound was dispensed directly into the wells via acoustic transfer (Beckman, ECHO 550R series). After 2h of incubation (37°C, 5% CO_2_), plates were equilibrated to RT for 30 min. 10 μL of NanoBiT substrate (Promega, Nano-Glo Live Cell Reagent, N2011) was added to each well by Matrix WellMate dispenser (Small-Bore Nozzle, ThermoScientific 201- 30002) and plates were shaken for 5 minutes at 50 x *g* on an orbital shaker prior to incubation for 25 min at RT. Luminescence was measured using an Envision Multilabel reader (200 ms). Readings from all DMSO wells were averaged (DMSO = 100%) and each compound treated well was normalized to DMSO. Results were plotted in GraphPad PRISM 9.5.1 and data were fit using the non-linear fit module (4- parameter - log agonist versus response).

### ProLabel protein degradation assay

#### NEK7

NEK7 was cloned into pLenti6.2-V5-Prolabel using standard gateway protocols (Gateway LR Clonase II Enzyme mix, 11791020). All DNA inserts were sequence verified. 293T and 293GT CRBN KO were transfected as follows: on day 1 cells were diluted in (DMEM) growth media to 1×10^6^ and plated into 6- well dishes (Corning, 3516), 2 mL/well, and then incubated overnight (37° C, 5% CO_2_). On day two, Fugene HD Transfection Reagent (Promega, E2311) and DNA were diluted into 200 µL OptiMEM^TM^ (ThermoFisher Scientific, 3185062) at a ratio of 3:1 Fugene to DNA, 3 µg of DNA per well. Mixture was incubated at RT for 25min and then added dropwise to cells, 200 µL/well, and then incubated overnight (37° C, 5% CO_2_). On day 3 cells were trypsinized (ThermoFisher Scientific, TrypLE Express Enzyme, 12605010), counted and diluted in DMEM growth media to 5×10^5^ cells/mL, plated at 20 µL/well in solid white 384 well plates (Corning 3570), and incubated overnight (37° C, 5% CO_2_). Test compounds were serially diluted in DMSO and 100 nL of compound was dispensed directly into the wells via acoustic transfer (Beckman, ECHO 550R series). After 18 h of incubation (37°C, 5% CO_2_), plates were equilibrated to RT for 1h. To detect Prolabel-NEK7 levels, 15 µL of DiscoverX PathHunter Prolabel Detection Reagent (User Manual 93-0180) was added to each well by Matrix WellMate dispenser (Small-Bore Nozzle, ThermoScientific 201-30002) and plates were shaken for 5min at 50 x *g* on an orbital shaker prior to incubation for 3h at RT. Luminescence was measured using an Envision plate reader (200 ms Luminescence, luminescence mirror module 2100-404, Emission filter 212) and luminescence values for each test compound-treated well was normalized to the average values for all DMSO wells within a given plate. Readings from all DMSO wells were averaged (DMSO = 100%) and each compound treated well was normalized to DMSO. Results were plotted in GraphPad PRISM 9.5.1 and data were fit using the non-linear fit module (4-parameter - log agonist versus response).

#### IKZF2

To measured IKZF2 degradation, HEK293Tcells stably expressing Prolabel-IKZF2 were used. Cells were trypsinized and diluted in DMEM growth media to 5×10^5^ cells/mL plated at 20 µL/well in solid white 384 well plates (Corning 3570), and incubated overnight (37°C, 5% CO_2_). Test compounds were serially diluted in DMSO and 100 nL of compound was dispensed directly into the wells via acoustic transfer (Beckman, ECHO 550R series). After 18h of incubation (37°C, 5% CO_2_), plates were equilibrated to RT for 1h and Prolabel-IKZF2 levels were detected and plotted as described above for NEK7.

### Molecular Modelling

The NEK7 beta-hairpin loop RMSD to CK1a beta-hairpin loop was determined to be 0.25 Å using the Biopython Superimposer module ^50^ and carbon-α atoms only (NEK7 PDB: 2WQN ^10^, where the loop is ^51^AACLLDGVPV^60^; CK1a PDB: 5FQD ^27^, where the loop is ^34^AINITNGEEV^43^).

The CRBN V338I point mutation was introduced into our cryo-EM structure of CRBN-NK7-902-NEK7 using CCG MOE *(Molecular Operating Environment (MOE)*, 2022.02 Chemical Computing Group ULC, 910-1010 Sherbrooke St. W., Montreal, QC H3A 2R7, 2024) and the side-chain minimized using the protein builder module. Illustrative figures were generated using PyMOL (*The PyMOL Molecular Graphics System,* Version 3.0, Schrödinger, LLC).

### Protein expression and purification

Both human NEK7(1-302) and mouse NEK7(1-302) were subcloned into a BL21-derived vector backbone with a N-terminal His6 tag followed by a ZZ tag and human rhinovirus 3C protease cleavage site. For the human NEK7 an additional 3x(GGGS) linker was inserted between the ZZ tag and the cleavage site. *E. coli* BL21(DE3) cells, transformed with the respective expression plasmid were cultivated shaker flasks containing Terrific Broth modified medium supplemented with MOPS buffer (Avantor) and Kanamycin (Avantor, K0047-5G). After 2h of incubation at 37°C and an OD_600_ value of 1.5 the culture was cooled down to 18°C. After the cultured reached an OD_600_ value of 3.5 expression was induced by addition of IPTG (Avantor.11IPTG0005) to a final concentration of 0.5 mM. Protein expression was performed overnight and cells were harvested by centrifugation and frozen at −20°C.

For purification, the cell pellet was resuspended in buffer A (50 mM HEPES, pH 7.5, 500 mM NaCl, 30 mM imidazole, 1 mM tris(2-carboxyethyl)phosphine, 5% glycerol; supplemented with 10 units/mL Benzonase nuclease (Sigma-Aldrich, 70746-4), and cOmplete, EDTA-free Protease Inhibitor Cocktail (Sigma-Aldrich; 4693132001, one tablet for 100 mL buffer) and ruptured by sonication. After centrifugation and filtration, the supernatant was loaded on a gravity flow column loaded with Talon Resin (Takara Bio, 635504). This and all subsequent purification steps were performed at 4°C. The target protein was eluted in buffer A which was supplemented with 500 mM Imidazole. Next, the His-tag was proteolytically removed by incubating the samples in the presence of 1:80 (w/w) human rhinovirus 3C protease (in-house produced) for 12h at 4°C. Optional dephosphorylation was performed by the addition of Lambda phosphatase (in-house) for 12-14h at 4°C with complete dephosphorylation confirmed by intact -LC-MS analysis.

After concentrating the samples in an Amicon Ultracentrifugation device (30 kDa cutoff), 5 mL of the samples were applied to a size exclusion chromatography column (Superdex 200pg, HiLoad 16/60, GE healthcare) equilibrated in SEC buffer (20 mM HEPES, pH 7.5, 300 mM NaCl, 0.5 mM tris(2- carboxyethyl)phosphine, 5% glycerol). Fractions containing the desired protein were pooled and concentrated in an Amicon Ultracentrifugation device (30 kDa cutoff). The concentrated protein was flash frozen as 100 μL aliquots in liquid nitrogen and stored at −80°C.

Biotinylated, Avi-tagged full length human CRBN in complex with human, full length DDB1, full length human His-PreSc-CRBN(1-442) in complex with full length human DDB1(1-1140) as well as mouse, full length CRBN in complex with mouse DDB1 were produced as previously described ^28^.

### SPR CRBN binding measurements

Binding of compounds to CRBN was assessed using SPR by immobilizing biotinylated DDB1:N-avi-CRBN complex (immobilized to ∼8000 RUs) on a Biacore streptavidin sensor chip and then injecting a dilution series of each compound. Experiments were performed with a Biacore T200 instrument with a 12 point, 2-fold dilution series and a variable top concentration, typically between 1 - 10 μM compound. All experiments were conducted at a temperature of 15°C with a flow rate of 30 µL/minute in phosphate-buffered saline (PBS) buffer at pH 7.2 containing 150 mM NaCl (300mM total NaCl), 2 mM EDTA, 5% glycerol, 0.01% Tween-20, 2% DMSO, and 1 mM TCEP. Data analysis to determine kinetic and steady-state affinity was performed using the Biacore Insight Evaluation Software.

### SPR assessment of ternary complex formation

Ligand affinities in the ternary complex (KD) were determined by SPR using a BiacoreTM T200 device (GE Healthcare). Biotinylated, Avi-tagged human and mouse CRBN/DDB1 complexes were immobilized on a Streptavidin-coated sensor (Cytiva, BR-1005-31) to a density of ∼5000 RU. The running buffer contained 20 mM TRIS-HCl pH 8, 200 mM NaCl, 1 mM TCEP, 0.05% Tween-20, 5% glycerol and 1% DMSO.

Experiments were carried out at 15°C using a flow-rate of 35 µL/min. The buffer of human and mouse NEK7 was exchanged on Amicon Ultracentrifugation devices (30 kDa cutoff) to the SPR running buffer. Varying concentrations of these proteins were mixed with a fixed concentration of molecular glue at 50 µM.

These mixtures were tested in standard successive injection mode at nine different concentrations. Curve fitting was performed using the Biacore T200 evaluation software using a steady state fit and GraphPad Prism.

### EM sample preparation and data collection

hNEK7 (1-302) was concentrated with a Amicon Ultracentrifugation device (30 kDa cutoff, Merck) to 25.42 mg/mL. Human His-PreSc-CRBN(1-442)//DDB1(1-1140) complex at 10.67 mg/mL in 50mM HEPES pH 8.0, 200mM KCl, 0.25 mM TCEP (uncleaved) was mixed with hNEK7 (1-302) at 25.42 mg/mL, NK7- 902 (1 mM in 10% DMSO) and buffer containing 50mM HEPES pH 7.5, 150mM NaCl (freshly filtered through 0.22 µm filter, Merck) resulting in a mixture of final 6 µM CRBN/DDB1, 50 µM hNEK7 and 100 µM NK7-902. Samples were incubated for 20min on ice and centrifuged for 10min at 16,000 x *g* and 4 °C.

This solution was used for cryo-EM grid preparation. Quantifoil gold grids (Au 200, R1.2/1.3, Quantifoil) were glow discharged for 60sec at 20 mA using Glocube (GloQube, Quorum). Five μL of sample were applied to a glow-discharged grid. The grid was plunged in liquid ethane after blotting for 5sec using blot force of 25 in 4 ℃ temperature and 100% humidity. Plunging was done using a Vitrobot Mark IV system (ThermoFisher Scientific),

Data collection was performed on a Glacios transmission electron microscope (Thermo Fisher Scientific) operated at 200 kV. EPU software was used to collect the dataset at a nominal magnification of 120 K and spot size 4, resulting in a physical pixel size of 0.845 Å using F4i camera (ThermoFisher Scientific). The data was acquired using the dose rate of 15.99 e- /Å2S, and a total dose of 50.21 e- /Å2. Each movie was fractionated into 50 subframes. The images were recorded at a defocus range of −0.8 to −1.6 μm and 0.2 μm increment.

### Image processing and 3D reconstruction

25833 micrographs were motion corrected using MotionCor2 ^51^. Non-dose weighted micrographs were used for Ctf-estimation using Gctf ^51^. Dose-weighted micrographs were used to pick particles in CisTEM ^52^. Particles were extracted using 300 px box. A homogeneous subset of particles was obtained after two rounds of 2D – classification in CisTEM. 455956 particles from cisTEM were imported and extracted in Relion ^53^. Extracted particles in Relion were imported in CryoSPARC ^54^. Two rounds of 2D – classification and ab-initio were done to remove the particles where NEK7 density was not observed. As a result, 327074 particles were used to do non-uniform refinement in CryoSPARC. Finally, the last clean subset of particles yielded a final map with a global resolution of 3.4 Å based on the gold-standard 0.143 criterion.

### Model building

Deposited models of hCRBN/DDB1-CK1a complex (pdb ID: 5FQD) and a Alphafold2 structure of hNEK7 (AF-Q8TDX7-F1) were docked in the map using Chimera ^55^. Manual model building was performed in Coot ^56^ and real space refinement was carried out in Phenix ^57^. Validity of the structure coordinates was assessed by Molprobity ^58^. The coordinates and EM-maps have been deposited under the following accession code: PDB ID 9H59, EMD-51881.

### Human subjects

For *in vitro* studies, fresh human blood was collected from healthy donors, from which no additional details are available, with informed consent (Santé med Gesundheitszentrum AG Basel, Switzerland). Buffy coats were obtained from healthy donors, from which no additional details are available, from Interregionale Blutspende (IRB) SRK Bern.

Blood from CAPS patients was collected at the University Children’s Hospital Tübingen, with approval for use of biological material obtained by the local ethics committee of the Medical Faculty Tübingen (approval 1015/2020BO2), and written informed consent obtained before blood drawing in accordance with the Declaration of Helsinki and local legislations. Blood samples were further processed in the lab of Prof. Weber as described in the next section. The below information was provided for the different patients:

| Patient ID | NLRP3 mutation | Gender | Age (years) | Treatment |
| --- | --- | --- | --- | --- |
| 1 | H360A | male | 17 | Canakinumab |
| 2 | A439M | male | 27 | Canakinumab |
| 3 | T348M | male | 40 | Canakinumab |
| 4 | T348M | female | 60 | Canakinumab |
| 5 | E311K | female | 32 | Canakinumab |
| 6 | E311K | female | 53 | Canakinumab |
| 7 | E311K | female | 60 | Canakinumab |

### Cells and culturing conditions

#### Human THP1 cells

Human THP1 cells (ATCC-TIB-202) were cultured in RPMI 1640 containing Glutamax (GIBCO, 72400- 021) and 25 mM Hepes, supplemented with 10% heat inactivated (HI) FBS, 1% Penicillin/Streptomycin, 1 mM Sodium Pyruvate and 50 µM β-mercaptoethanol.

#### Human iPS cells

Female human WT29 iPS cells were derived from cell line AG092429 obtained from the NIA Aging Cell Repository at the Coriell Institute for Medical Research. Stem cells were plated on a LN511-coated surface (BioLamina LN521) and amplified in mTeSR Plus medium (Stemcells Technologies, 05825) supplemented with penicillin/streptomycin (GIBCO, 15140). During the first 24h after seeding, the medium was supplemented with 10 µM ROCK Inhibitor (Y-27632) (Stemcells Technologies, 72305). The medium was changed every 24h and the cells were passaged every 3 days. For passaging, the cells were washed with PBS (GIBCO 14190-094) and detached by incubation with TrypLE (GIBCO, 12604-013) at 37°C for 5 min. EB formation was initiated by putting 3×10^6^ cells on a 6-well AggreWell (Stemcells Technologies, 34811) according to the manufacturer’s protocol in mTeSR PLUS was supplemented with penicillin/streptomycin and 10 µM ROCK inhibitor (Y-27632 Tocris, 1254). Three days later about 200 EB were transferred to a well of a 24-well ultra-low attachment plate (Corning, 3473) and treated with the compounds. For Proteomics, the EB’s were washed twice with PBS and frozen at −80°C.

#### Human PBMCs and monocytes

Human PBMCs were isolated from fresh whole blood collected in EDTA tubes at RT. diluted 1:3 in PBS and layered onto Leucosep tubes containing 15 mL Ficoll (Greiner Bio-One, 227289). Tubes were centrifuged at 1000 x *g* for 20min, without brake, at RT. The PBMC layers were collected and washed with Facs Buffer (PBS,1% HI-FBS, 2mM EDTA). Supernatant was removed by aspiration, PBMCs were resuspended with Facs Buffer and centrifuged at 200 x *g* for 10 min, with low break, at RT to eliminate platelets. Erythrocytes were lysed by resuspending the cell pellet in 10 mL Qiagen EL Buffer (Qiagen, 79217), followed by a 5min incubation at RT. Tubes were filled up with Facs Buffer to 50 mL total volume and centrifuged at 500 x *g* for 5min before being re-suspended in complete medium and counted.

Human monocytes were isolated from human PBMCs using the EasySep Human Monocyte isolation kit (Stem Cell, 19669), following the manufacturer’s instructions. Human PBMCs and monocytes were cultured in RPMI 1640 containing Glutamax and 25 mM Hepes, supplemented with 10% HI-FBS, 1 mM sodium pyruvate, 1% penicillin/streptomycin and 50 µM β-mercaptoethanol

#### Human CAPS PBMCs

PBMCs from CAPS patients were prepared as follows at the Department of Innate Immunity of the University of Tübingen. Whole blood was collected in EDTA tubes at RT. Samples were processed within 30min upon collection. Plasma was separated by centrifuging acquisition tubes at 700 x *g* for 5min at RT. Plasma was transfer to sterile 15 ml falcons. From the remainder, PBMCs were isolated by density-gradient centrifugation using Histopaque-1077 Separation Solution (Sigma-Aldrich). Whole blood was first diluted with two times the volume of PBS and 22 mL aliquots of the blood solution were then layered onto 18 mL Ficoll in 50 mL centrifugation tubes. The tubes were centrifuged at 800 x *g* for 20min, acceleration 4, without brake and at RT. The PBMC layers were collected and washed with PBS and a centrifugation step at 650 x *g* for 8min; the resulting pellets were resuspended in 15 mL PBS, all cell suspensions were pooled in one 50 mL Falcon tube and centrifuged at 450 x *g* for 8min. PBS was completely removed by vacuum aspiration. Erythrocytes were lysed by resuspending the cell pellet in 1 mL ACK lysing buffer (ThermoFisher Scientific, A1049201), followed by a 4min incubation at RT. The sample was then filled up with PBS to 20 mL total volume. Centrifugation was carried out at 250 x *g* for 8min. The resulting cell pellet was re-suspended in 5 ml complete medium and counted. Subsequently the cells were frozen as follows: centrifugation at 500 x g for 5min, removal of the supernatant by pouring out, and resuspending the cells in freezing media (90% HI-FBS, 10% DMSO) at RT. 1 mL cell aliquots were transferred to 2 mL labelled cryovials (10-15×10^6^ cells per vial in 1 ml aliquots). The cryovials were frozen at −80°C overnight using a Coolcell LX Cell Freezing Container. On the next day, the frozen vials were transferred to liquid nitrogen for subsequent shipment to the Novartis research center. Aliquots of frozen PBMCs from healthy donors (prepared at Novartis following the same procedure described for CAPS PBMCs) were used as controls.

#### Human neutrophils

Human neutrophils were isolated from whole blood using the EasySep Human Neutrophils isolation kit (Stem Cell, 19666), following the manufacturer’s instructions. Once isolated, neutrophils were cultured in RPMI 1640 Glutamax, 10.000 UL penicillin/streptomycin, aprotinin 15 mg/mL (Sigma; 3428) and 2% HI autologous plasma. The latter was obtained as follows: 4 mL of undiluted blood were layered on 2 mL of Ficoll Paque Plus (Cytiva, 17144003) and tubes were spun at 2200 rpm for 20min, breaks off. One to two mL of plasma was collected and heat inactivated in a water bath at 56°C for 30min. Plasma was then cooled down, centrifuged for 10min at 1500 x *g* to remove any debris and finally filtered through a 0.22 µM filter before being added to the culture media.

#### Mouse primary splenocytes

For the isolation of mouse primary splenocytes, spleens were collected in PBS from naïve C57BL/6 wild type mice and mashed through a 70 µM cell strainer using the plunger of a 5 mL syringe. Cells were washed with Facs Buffer (PBS,1% HI-FBS, 2mM EDTA) and incubated for 10min at RT with 10 mL of red blood cell lysis buffer (Qiagen). Cells were washed again with Facs buffer, filtered through a 40 µM cell strainer and resuspended in complete medium and counted. All cells were cultured in a humidified incubator at 37°C and 5% CO_2_.

### WES capillary electrophoresis immunoanalysis

Human and mouse primary cells were collected in eppendorf tubes, centrifuged at 500 x *g* for 5min at 4°C, washed with ice-cold PBS and centrifuged again at 700 x *g* for 5min at 4°C. Cell pellets were resuspended in 20-100 µL of RIPA buffer (Thermo Fisher Scientific, 89901) containing 1x protease and phosphatase inhibitors (Thermo Fisher Scientific,78444), vortexed and transferred to −80°C overnight to complete the lysis.

Human and NHP whole blood samples were processed as follows: 200 µL of blood were diluted with 200 µL of complete medium (RPMI 1640 Glutamax, 5% heat inactivated FCS, 10 mM Hepes, 50 µM β-mercaptoethanol) and centrifuged at 500 x *g* for 10min at RT. The upper fraction containing platelets was removed and red blood cells were lysed with 1.5 mL 1x ACK buffer diluted in water (10x ACK buffer: 1.5 M NH_4_Cl, 100 mM KHCO_3_, 0.99 mM EDTA) for 15min on ice. Samples were spun down at 300 x *g* for 5min at 4°C, white blood cells were washed twice with ice-cold PBS and resuspended with 60-80 µL of RIPA buffer containing 1x protease and phosphatase inhibitors and 5 µM diisopropylfluorophosphate (Sigma D0879). Samples were vortexed and incubated overnight at −80°C to complete the lysis.

Mouse whole blood samples were processed as follows: 200 µL of blood was incubated with 2ml of 1x Red Blood Cell Lysis buffer diluted in water (Miltenyi Biotec #130-094-183) for 10min at RT. Samples were spun down at 300 x *g* for 5min. White blood cells pellets were washed with PBS and resuspended with 40 µL of RIPA buffer containing 1x protease and phosphatase inhibitors. Samples were vortexed and incubated overnight at −80°C to complete the lysis.

The next day, primary cell and whole blood lysates were thawed on ice and centrifuged at 4100 x *g* for 30min at 4°C. The supernatant was collected and transferred to a 96 microwell plate. Total protein concentration was determined for each sample using the Pierce BCA assay (Thermo Fisher Scientifc 23225). Cell lysates were then prepared for automated Western blot (WES/Jess) following instructions and using reagents provided by Protein Simple. Briefly, 5 µL of sample containing 2 µg of denatured protein mixed with fluorescent dye were loaded in a 25-well microplate. 10 µL of anti-NEK7 (Abcam, 133514, 1:500) and anti-actin antibodies (Cell Signaling Technology, 1:200) and 10 µL of secondary anti-rabbit antibody (ready to use, Protein Simple) were loaded in the appropriate wells. All assay steps including separation, immunodetection and data analysis were performed by the WES/Jess instruments. Data were analyzed and plotted using the Compass for Simple Western and GraphPad Prism software.

### Cytokine assays *in vitro*

Human monocytes and PBMCs were plated in 384-well plates at the concentration of 17’000 cells/well and 33’000 cells/well, respectively, pre-incubated with compounds and stimulated with either LPS (LPS- EB Ultrapure, Invivogen tlrl-3pelps) alone, LPS plus ATP (Sigma, A2383), LPS plus nigericin (Enzo Life Sciences, BML-CA421-0005) or LPS plus niclosamide (Adipogen, AG-CR1-3643-M100) as indicated in the figure legends. Human and NHP blood was diluted 1:2 in whole blood medium (RPMI + GlutaMax, 5% HI-FBS, 10mM HEPES, 50uM β-mercaptoethanol), plated in 384-well plates (43ul/well), treated with compounds and stimulated with LPS and ATP as indicated in the figure legends. At the end of the stimulation, monocytes and PBMCs plates were centrifuged at 500 x *g* for 5 min while whole blood plates were centrifuged at 200 x *g* for 2min, supernatants were collected and IL-1β levels were measured using the Human IL-1β HTRF kit (Cisbio – PerkinElmer), following the manufacturer’s instructions. Plates were read with a PHERAstarFSX instrument.

Human neutrophils were plated in 96-well plates at the concentration of 200’000 cells/well incubated with the indicated concentrations of NK7-902 or NP3-253 for 18h and then stimulated LPS plus ATP as indicated in the figure legends. IL-1β levels in the supernatant were measured using U-plex Biomarker Group1 human kit (Meso Scale Discovery), following the manufacturer’s instructions.

### Expression proteomics

One to two million cells were treated overnight with DMSO or 1 µM NK7-902, washed twice with PBS and then stored as dry pellets at −80°C until further processing. Each condition was tested at least in duplicate. TMT-labeled peptides were generated with the iST-NHS kit (PreOmics) and the TMT11plex or TMT16plex reagent (Thermo Fisher Scientific). Equal amounts of labeled peptides were pooled and separated on a high pH fractionation system with a water-acetonitrile gradient containing 10 mM ammonium formate, pH 11. Alternating rows of the resulting 72 fractions were pooled into 24 samples, dried and resuspended in water containing 0.1% formic acid. The LC-MS analysis was carried out on an EASY-nLC 1200 system coupled to an Orbitrap Fusion Lumos Tribrid mass spectrometer (Thermo Fisher Scientific). Peptides were separated on a 25 cm long Aurora Series UHPLC column (Ion Opticks) with 75 µm inner diameter and a water-acetonitrile gradient containing 0.1% formic acid over 180 min. MS1 spectra were acquired at 120k resolution in the Orbitrap, MS2 spectra were acquired after CID activation in the ion trap and MS3 spectra were acquired after HCD activation with a synchronous precursor selection approach using five notches and 50k resolution in the Orbitrap. LC-MS raw files were analyzed with Proteome Discoverer 2.4 (Thermo Fisher Scientific). Briefly, spectra were searched with Sequest HT against the Homo sapiens UniProt protein database and common contaminants (2019, 21494 entries) or Mus musculus and common contaminants (2019, 23082 entries). The database search criteria included: 10 ppm precursor mass tolerance, 0.6 Da fragment mass tolerance, maximum three missed cleavage sites, dynamic modification of 15.995 Da for methionines, static modifications of 113.084 Da for cysteines and 229.163 Da or 304.207 Da for peptide N-termini and lysines. The Percolator algorithm was applied to the Sequest HT results. The peptide false discovery rate and the protein false discovery rate was set to 1%. TMT reporter ions of the MS3 spectra were integrated with a 20 ppm tolerance and the reporter ion intensities were used for quantification. The mass spectrometry proteomics data have been deposited to the ProteomeXchange Consortium via the PRIDE ^59^ partner and reviewer access will be provided upon request.

Protein relative quantification was performed using an in-house developed R script, available on GitHub (https://github.com/Novartis/px_tmt_daa). This analysis included multiple steps; (1) data filtering (exclusion of peptides mapping to multiple proteins, exclusion of PSMs where the number of SPS mass matches were <60%, the precursor interference was >50% or the average reporter ion s/n <10, as well as exclusion of PSMs with missing reporter ion signals); (2) global data normalization by equalizing the total reporter ion intensities across all channels, (3) summation of reporter ion intensities per protein and channel, calculation of protein abundance log2 fold changes (L2FC) and testing for differential abundance using moderated t-statistics ^60^ where the resulting p values reflect the probability of detecting a given L2FC across sample conditions by chance alone. Subsequently, the p values were adjusted for multiple testing using the Benjamini–Hochberg method (q values).

### Generation of NEK7 KO cells and cell cycle analysis

HeLa and U2OS cells were confirmed by SNP analysis and verified to be mycoplasma-free by routine testing. All cells were kept in a humidified incubator at 37°C and 5% CO_2_ and were maintained in DMEM supplemented with 10% FBS, 1% L-glutamine and 1% Penicillin/Streptomycin. Cell culture reagents were obtained from Invitrogen. HeLa and U2OS Cas9 cells were generated by lentiviral delivery of Cas9 in pNGx-LV-c004 followed by selection with blasticidin (PMID: 27351204). Control and NEK7 KO cells were generated by lentiviral delivery of non-targeting or NEK7 sgRNAs (TGGTACTCCATCCAAGAGAC) in pNGx-LV-g003 followed by selection with 1 μg/mL puromycin (PMID: 27351204). KO polyclonal cell populations were analyzed 7 days post-infection with sgRNA for efficiency of knockout by western blotting.

Synchronization and cell cycle analysis was done as described in PMID: 28539406. Cells were treated with 100 ng/mL nocodazole (EMD Millipore, 487928) for 16h and released in fresh medium. Before flow cytometry, cells cultured on dishes for the indicated time were washed twice with PBS and fixed in 70% cold ethanol at −20°C for 3 h. The fixed cells were then washed with PBS and incubated with Muse Cell Cycle Reagent at room temperature for 30 min. The DNA contents of the cells were then measured using Muse Cell Analyzer (Merck Millipore). Parallel samples were analyzed by western blotting using the following antibodies at 1:1000 dilution: rabbit polyclonal antibodies against STIL (ab89314; Abcam), NEK7 (3057S; Cell Signaling Technology), CPAP/CENP-J (11517-1-AP; Proteintech), GAPDH (2118S; Cell Signaling Technology); mouse monoclonal antibodies against p21 Waf1/Cip1 (DCS60; Cell Signaling Technology). The CDK and cyclin antibodies were used from the CDK and Cyclin Antibody Sampler Kits (9868 and 9869, respectively; Cell Signaling Technology).

### Micronucleus test in cultured human peripheral blood lymphocytes

Heparinized blood from healthy volunteers was used. 5 mL cultures of 0.4 mL whole blood in RPMI medium containing 20% fetal bovine serum, 1% penicillin/streptomycin and 37.5 μl/mL phytohaemagglutinin (PHA) were cultivated at 37°C. Cytochalasin B (6 µg/mL) was added 72h after inoculation and cultures were harvested at 96h. Cells underwent cold hypotonic treatment (0.075 M KCl) and were immediately centrifuged and fixed three times with fixative (methanol:acetic acid, 3:1) supplemented with 1% formaldehyde for the first fixation. The fixed cells were dropped onto clean microscope slides (two slides per culture), air-dried, and stained with 2% Giemsa in a 5-mM Na2HPO4–5- mM NaH2PO4 buffer and mounted in Terpex (LabForce AG, Nunningen, Switzerland). The test compound was first added at 44h after PHA stimulation. To allow continuous exposure to the test compound, cell cultures were washed and treated again with the test compound every 24h up to 96h. Carbendazim (MBC, CAS no. 10605-21-7) was purchased from Sigma–Aldrich Chemie (Steinheim, Germany) and dissolved in DMSO.

As a measure of cytotoxicity and cell cycle delay the relative division index was used:

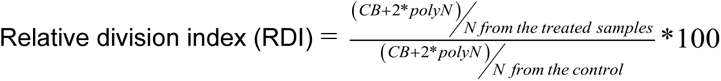

CB: Binucleated cells; PolyN: Polynucleated cells

The micronucleus test analysis was done using an image analysis system ROBIAS (Robotic Image Analysis System) for both automatic cytotoxicity assessment and micronucleus detection in human lymphocytes Six concentrations (two cultures per concentration) and 2000 binucleated cells per concentration were analyzed using a 100x oil immersion objective. A total of 500 cells/culture were scored to determine the division index DI (a cytotoxicity parameter) and 1000 binucleates analyzed for MN induction.

Statistically significant differences between controls and treated samples were determined with the chi-square test (95% confidence limit is used). A test compound was considered positive for the MN induction if it induced a reproducible statistically significant increase in the number of micronucleated binucleates (MNCB).

### Mice

All animal experimental procedures complied with the Swiss animal welfare regulations and were approved by the Cantonal Veterinary Office of Basel-City, Switzerland. Mice were housed under standard conditions with a 12-hour light/dark cycle (artificial light), a room temperature of 22°C (± 2°C) with a relative humidity of 55% (variable between 45 and 65%) and fed with standard chow and water provided ad libitum.

### Mouse peritonitis model

Female C57BL/6JRj mice (6 weeks old) were purchased from Janvier Labs (France) and randomized into 8 groups of 5 mice per cage. At an age of 7-8 weeks, C57BL/6JRj mice were dosed orally (10 mL/kg) either with vehicle (0.5% Methylcellulose (BioConcept 9-00F14-I), 0.1% Tween80 (VWR, M126) in 50 mM pH=4.7 acetate buffer (sodium acetate trihydrate, Sigma CAS# 6131-90-4; acetic acid, Sigma CAS# 64- 19-7 in water)) or NK7-902 at doses of 300, 100 and 30 mg/kg, in the same vehicle, 6h or 16h prior to priming of mice with an intraperitoneal (i.p.) injection of 2.5 mg/kg LPS (Lipopolysaccharide E. coli O111:B4, Sigma L2630, in 0.5 mL 1x PBS). Two hours later, the mice received an i.p. injection of 20 mM ATP (Adenosine triphosphate, Sigma A2383, in 0.5 mL 1x PBS) and 15 min thereafter, mice were euthanized with an overdose of CO_2_. After euthanasia, the peritoneal cavity was washed out using 3 mL of 0.9 % NaCl solution (VWR, 5929.1000) containing 1 mg/mL EDTA (VWR, M101), and the spleen was collected and snap frozen. The peritoneal washout fluid samples were centrifuged (250g, 10 min, 4 °C), and the cell-free supernatant was assayed for IL-1β concentrations using the mouse cytokine IL-1β set (Bio-Plex Pro mouse cytokine IL-1β set (Bio-Rad, 171G5002M), Bio-Plex Pro mouse cytokine standards group 1 (Bio-Rad, 171I50001), Bio-Plex Pro reagent kit V (12002798), according to manufacturer’s instructions.

### Mouse CAPS model

CAPS mice (mouse NLRP3^A350V^ Muckle-Wells Syndrome (MWS) model of CAPS) were bred internally by crossing B6.129-Nlrp3tm1Hhf/J mice (Prof. H. Hoffman, UCSD) with C57BL/6- Gt(ROSA)26Sortm9(Cre/ESR1)Arte mice (Taconic Biosciences), and heterozygous mice were then randomized into four groups of five mice per cage (two males and three females in each group). At an age of 14 weeks, mice were treated for five days with a daily dose of 5 mg/kg tamoxifen citrate salt > 99% (Sigma, T9262) by i.p. injection (10 mL/kg). Two days later, mice were dosed orally (10 mL/kg) either with vehicle (0.5% Methylcellulose (BioConcept 9-00F14-I), 0.1% Tween80 (VWR, M126) in 50 mM pH=4.7 acetate buffer (sodium acetate trihydrate, Sigma CAS# 6131-90-4; acetic acid, Sigma CAS# 64-19-7)), or NK7-902 at 300 mg/kg in the same vehicle, or NP3-253 at 20 mg/kg in vehicle (0.5% Methylcellulose (BioConcept 9-00F14-I), 0.1% Tween80 (VWR, M126) in water) 16h prior to priming of mice with an intraperitoneal (i.p.) injection of 10 µg/kg LPS (Lipopolysaccharide E. coli O111:B4, Sigma L2630, in 1x PBS, 10 mL/kg). Four hours later, mice were euthanized with an overdose of CO_2_ (>85%), blood from vena cava and the spleen were collected. Blood was processed to plasma and assayed for IL-1β concentrations using the mouse cytokine IL-1β set (Bio-Plex Pro mouse cytokine IL-1β set (Bio-Rad, 171G5002M), Bio-Plex Pro mouse cytokine standards group 1 (Bio-Rad, 171I50001), Bio-Plex Pro reagent kit V (12002798)), according to manufacturer’s instructions.

### NHP whole blood assay and PK/PD

Non-GLP single dose and multiple dose PK/PD studies with NK7-902 were conducted in male cynomolgus monkeys at Charles River Laboratories. The studies were conducted under the UK Home Office Project License No. PPL PP9376768 Drug Metabolism and Pharmacokinetics, Protocol Reference Number 3. The study protocols were approved by the local animal welfare and ethical review board (AWERB) prior to submitting to the Home Office. The study was conducted in compliance with the Animal Welfare Act, the Guide for the Care and Use of Laboratory Animals, and the Office of Laboratory Animal Welfare. The studies were carried out respecting all ethical regulations. Three male cynomolgus monkeys aged 3-4 years were administered orally 0.2 - 2 mg/kg/day NK7-902, resuspended in 20% (v/v) PEG300, 10% (v/v) Kolliphor (HS15), 70% (v/v) PBS pH 7.4, with final pH adjusted to pH 4-6 with 5 M HCl. Blood samples for PK were collected at the indicated time points by venipuncture in polypropylene tubes containing 1 M citric acid and frozen at −80°C until further analysis. Blood samples for PD (200 µl for measuring NEK7 degradation and 200 µl for measuring IL-1β inhibition following *ex-vivo* LPS stimulation) were collected in sodium heparin tubes at the indicated timepoints. Samples to measure NEK7 degradation were processed as follows: 200 µL of blood were diluted with 200 µL of complete medium (RPMI 1640 Glutamax, 5% heat inactivated FCS, 10 mM Hepes, 50 µM β-mercaptoethanol) were transferred to 2 mL eppendorf tube, centrifuged 500 x *g* 10min at RT. Upper fraction containing platelets was removed and red blood cells were lysed with 1.5mL 1x ACK buffer diluted in water (10x ACK buffer: 1.5 M NH_4_Cl, 100 mM KHCO_3_, 0.99 mM EDTA) 15min on ice and vortexed twice during lysis incubation. Samples were spun down 300g 5minutes at 4°C, white blood cells were washed twice with ice-cold PBS and centrifuged 700g 5min at 4C. Dry cell pellets were frozen on dry ice and stored at −80°C until further analysis. NEK7 protein levels were assessed as described in the “WES” section. To assess IL-1β inhibition, 200 µL of blood were stimulated in 96-well plates with LPS+ATP for 3.5h. Samples were spun down, serum was collected and frozen at −80°C until further analysis. Frozen serum samples were thawed on ice, diluted 1:40 in assay diluent 5 and then IL-1β levels were measured using the Human IL- 1β HTRF kit (Cisbio – PerkinElmer), following the manufacturer’s instructions. Plates were read with a PHERAstarFSX instrument.

### Data analysis

NEK7 recruitment and degradation as well as IL-1β inhibition data were analyzed and plotted using Microsoft Excel and GraphPad Prism.

## Supplemental Information

**Supplementary Figure 1.**
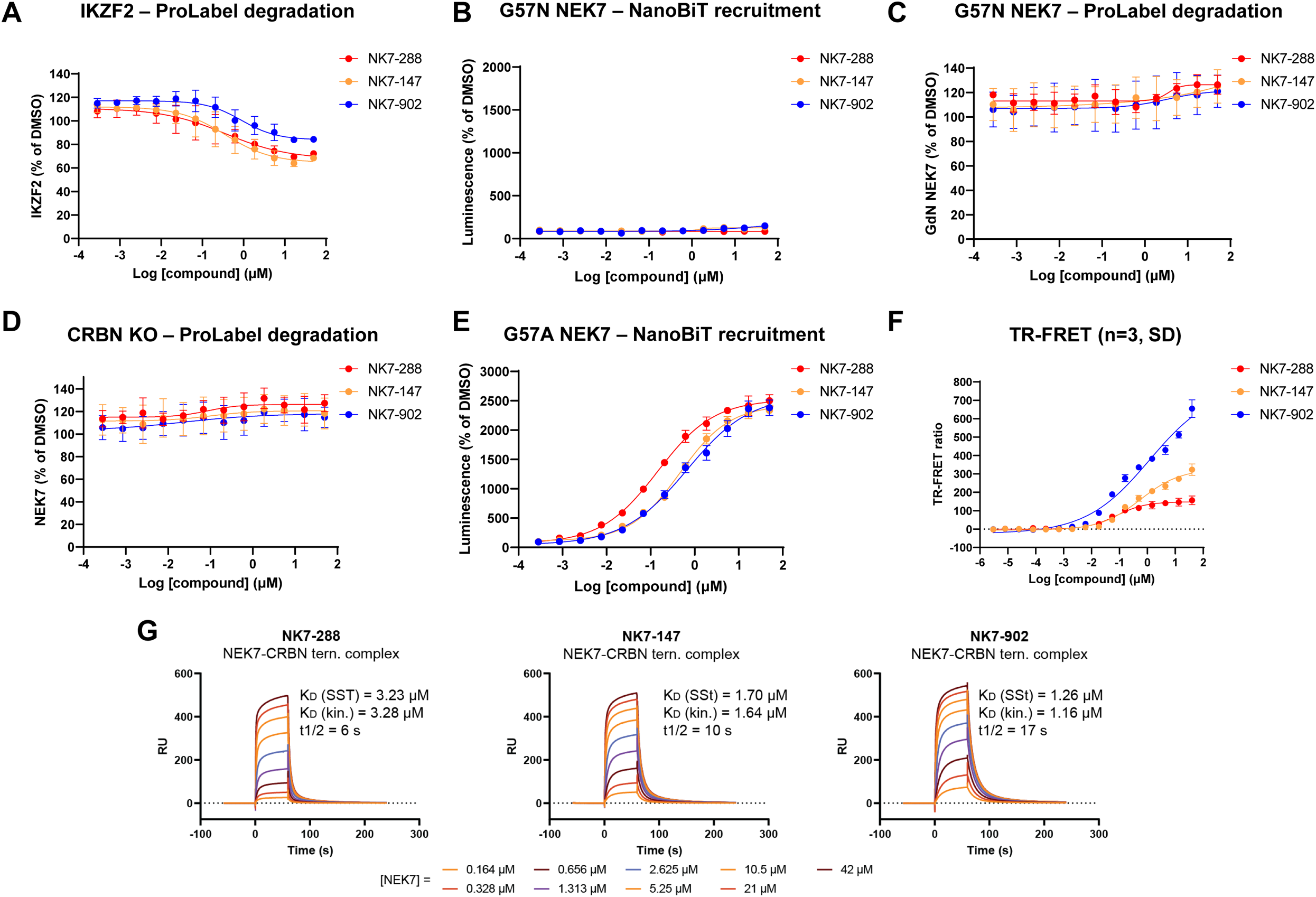
(A) 293T cells stably expressing ProLabel-tagged IKZF2 were incubated with increasing concentrations of NK7-288, NK7-147 or NK7-902 for 18h. Data represent mean ± SD (n=3). (B) 293T cells transiently transfected with plasmids encoding for SmBiT-tagged G57N NEK7 mutant and LgBiT-tagged CRBN were incubated with increasing concentrations of NK7-288, NK7-147 or NK7-902 for 2h in the presence of the NAE1 inhibitor MLN4924 as described in the methods. Data represent mean ± SD (n=8). (C) 293T cells were transiently transfected with a plasmid encoding for ProLabel-tagged G57N NEK7 and then incubated with increasing concentrations of NK7-288, NK7-147 or NK7-902 for 18h. Data represent mean ± SD (n=4) (D) CRBN KO 293T cells were transiently transfected with a plasmid encoding for ProLabel-tagged WT NEK7 and then incubated with increasing concentrations of NK7-288, NK7-147 or NK7-902 for 18h. Data represent mean ± SD (n=4). (E) 293T cells transiently transfected with plasmids encoding for SmBiT-tagged G57A NEK7 mutant and LgBiT-tagged CRBN were incubated with increasing concentrations of NK7-288, NK7-147 or NK7-902 for 2h in the presence of the NAE1 inhibitor MLN4924 as described in the methods. Data represent mean ± SD (n=8). (F) TR-FRET recruitment assay. Biotinylated NEK7 was incubated with a DDB1/GFP-tagged CRBN complex in the presence of increasing concentrations of NK7-288, NK7-147 or NK7-902 for 2min. Data represent mean ± SD (n=3). (G) SPR sensorgrams of the ternary complex of the immobilized CRBN/DDB1 complex with NEK7 and the respective compounds. Reported are the *K*_D_ values of a steady state fit (SSt) and kinetic fit (kin.) as well as the half-life of the ternary complex (*t*_1/2_).

**Supplementary Figure 2.**
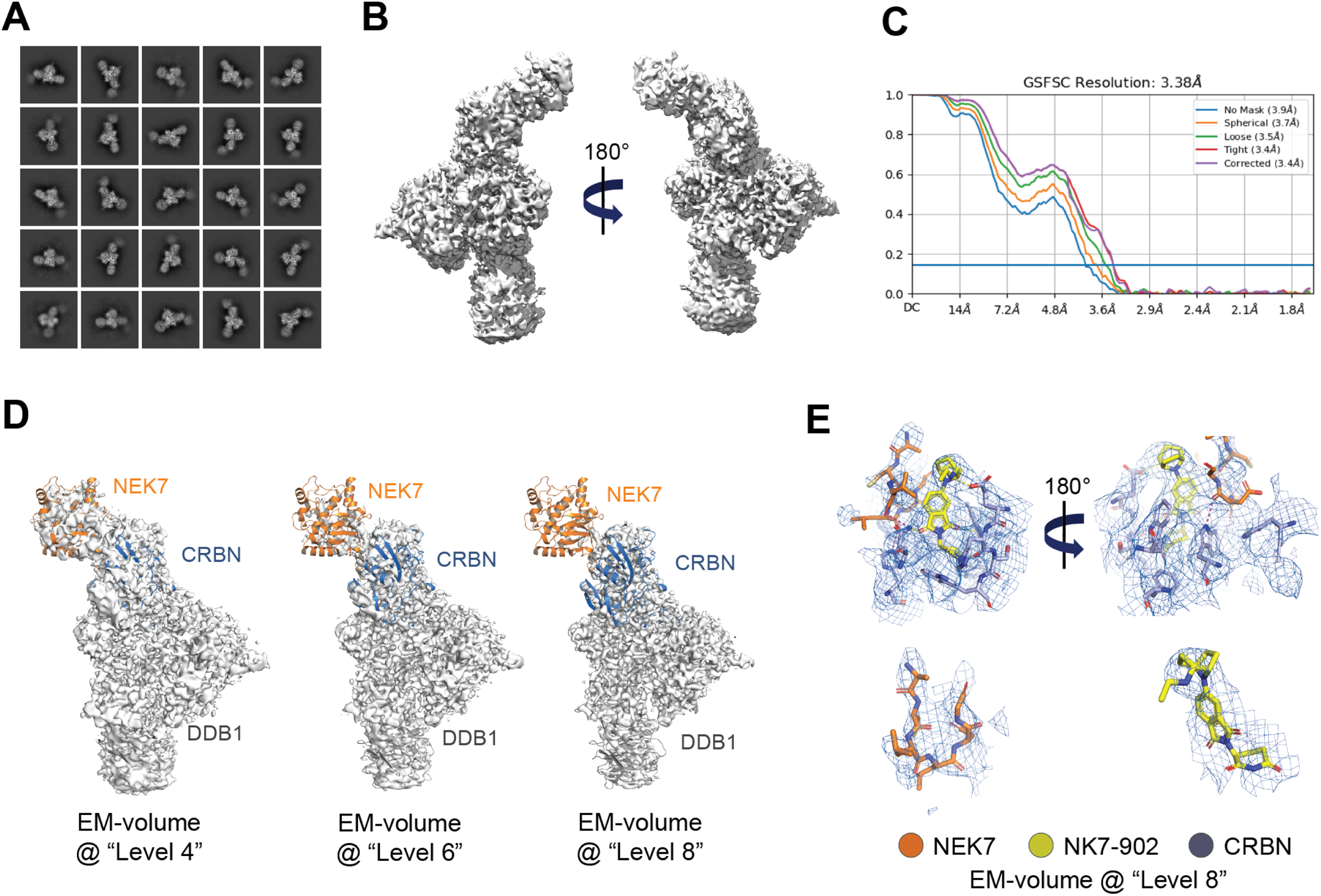
(A) Selected 2D classes of the cryo-EM data processing of the ternary complex of NEK7, NK7-902 and CRBN-DDB1. (B) Refined EM-map of the ternary complex of NEK7, NK7-902 and CRBN-DDB1. (C) FSC-based resolution analysis of EM-map of the ternary complex of NEK7, NK7-902 and CRBN-DDB1 indicating a maximal resolution of 3.38Å. (D) EM-volume of the ternary complex of NEK7, NK7-902 and CRBN-DDB1 at different scaling levels (as defined by Pymol) overlaid with the modeled ternary complex. (E) Modeled interactions of NEK7, CRBN and NK7-902 with the EM-map shown as blue dash. The same scaling level (“Level 8”) for the EM-map was used as in sub-panel (D).

**Supplementary Figure 3.**
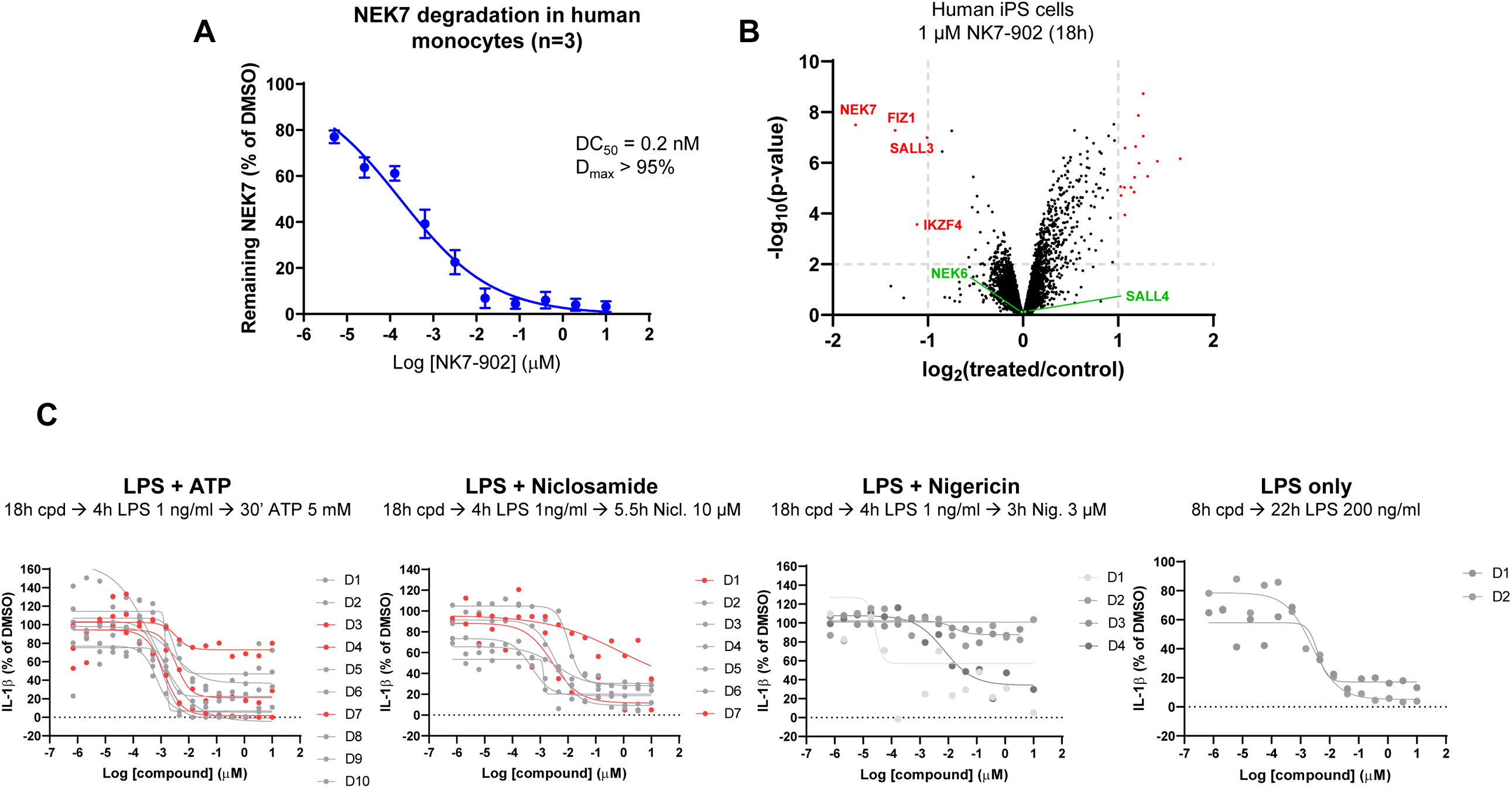
(A) Human primary monocytes were treated with increasing concentrations of NK7-902 for 18h and then lysed to measure NEK7 and β-actin levels by WES as shown in Figure 2A. For each sample, NEK7 levels were normalized over the corresponding β-actin signal. Data from three different donors were averaged and plotted ± SEM. (B) Human iPS cells were incubated with 1 µM NK7- 902 for 18h and then analyzed by LC-MS/MS to identify differentially modulated proteins. Proteins in the top left quadrant were significantly downregulated (see also Supplementary Table 2). (C) Chemical structure of the NLRP3 inhibitor NP3-253. (D) Individual curves for IL-1β inhibition data in monocyte shown in Figure 2D. Donors showing very different levels of IL-1β inhibition after activation with ATP or niclosamide are highlighted in red.

**Supplementary Figure 4.**
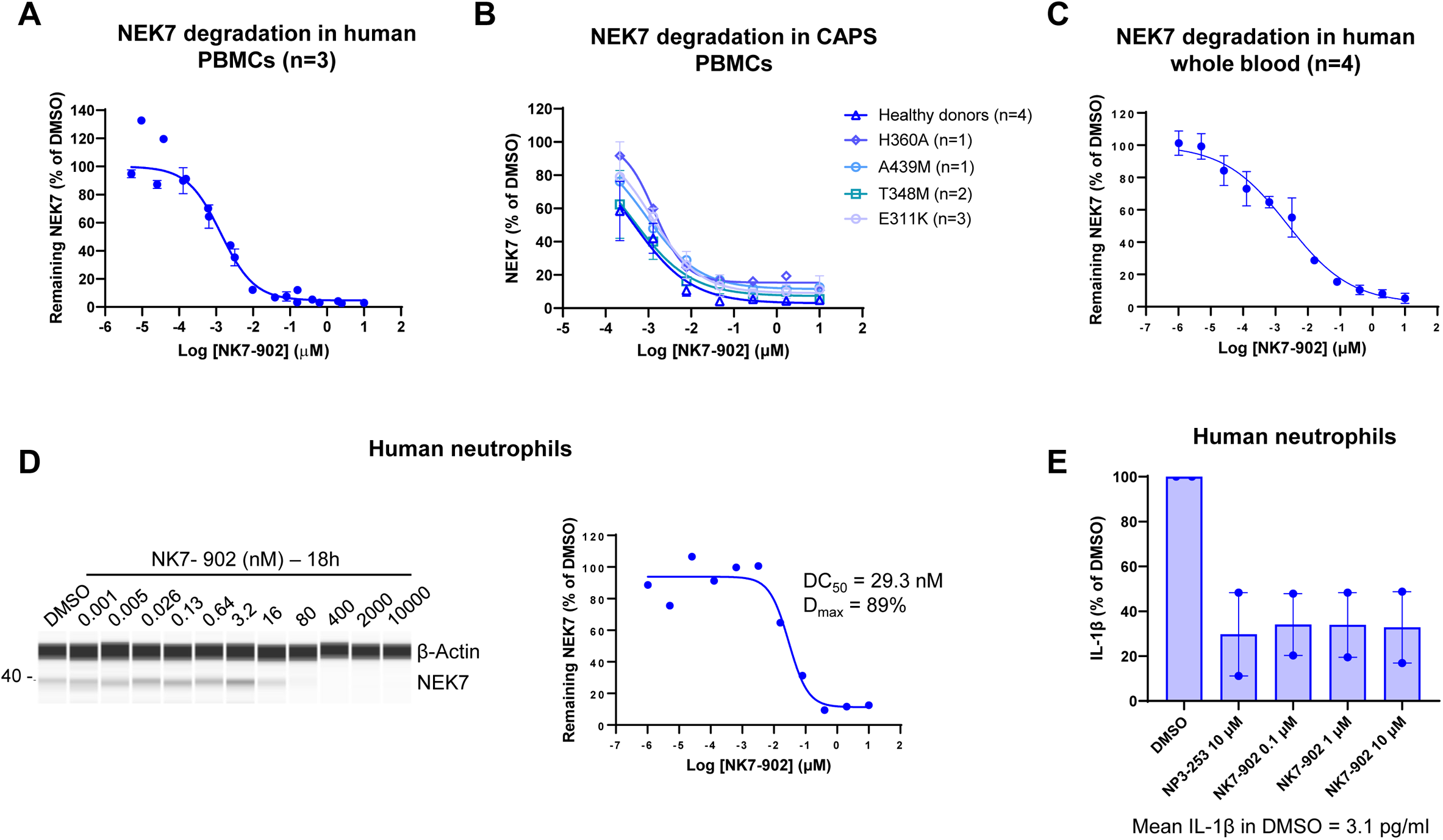
(A) Human PBMCs were treated with increasing concentrations of NK7-902 for 18h and then lysed to measure NEK7 and β-actin levels by WES as shown in Figure 3A. For each sample, NEK7 levels were normalized over the corresponding β-actin signal. Data from three different donors were averaged and plotted ± SEM. (B) PBMCs from healthy volunteers and CAPS patients carrying different NLRP3 mutations were incubated with increasing concentrations of NK7-902 for 18h and then lysed to measure NEK7 and β-actin levels by WES. For each sample, NEK7 levels were normalized over the corresponding β-actin signal. The number of donors tested for each NLRP3 mutation is indicated. Data represent mean ± SEM. (C) NEK7 degradation in human whole blood after treatment with NK7-902 was measured as described in Figure 3D. Data from four different donors were averaged and plotted ± SEM. (D) Human neutrophils were incubated with increasing concentrations of NK7-902 for 18h and then lysed to measure NEK7 and β-actin levels by WES. For each sample, NEK7 levels were normalized over the corresponding β-actin signal. Data were plotted on the left (n=1). (E) Human neutrophils were incubated with the indicated concentrations of NK7-902 or NP3-253 for 18h and then stimulated with 100 ng/mL LPS for 3h, followed by an additional 60min incubation with 3 mM ATP. IL-1β levels in the supernatant were measured by MSD. The mean IL-1β concentration (pg/mL) in the DMSO control is indicated below the graph (n=2).

**Supplementary Figure 5.**
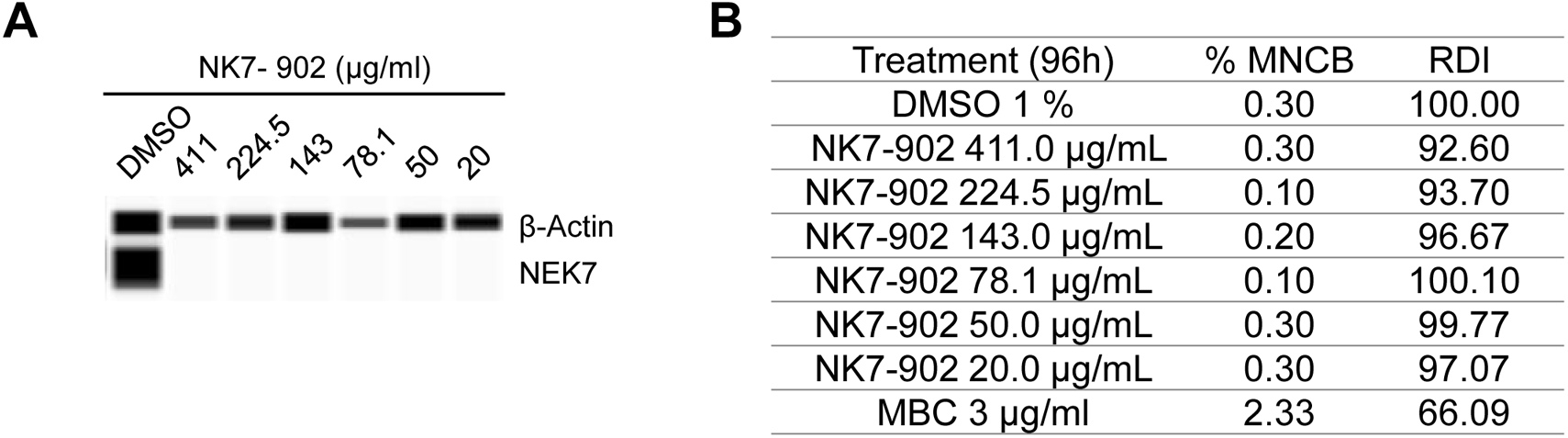
(A) Human whole blood lymphocyte cultures were incubated for 96h with the indicated concentrations of NK7-902 and then lysed to measure NEK7 and β-actin levels by WES. (B) Frequencies of micronucleated binucleates (MNCB) and the relative division indices (RDI) after incubation with the indicated concentrations of NK7-902. Carbendazim (MBC) was used as positive control. RDI: The relative division indices are used as a measure of cytotoxicity and cell cycle delay.

**Supplementary Figure 6.**
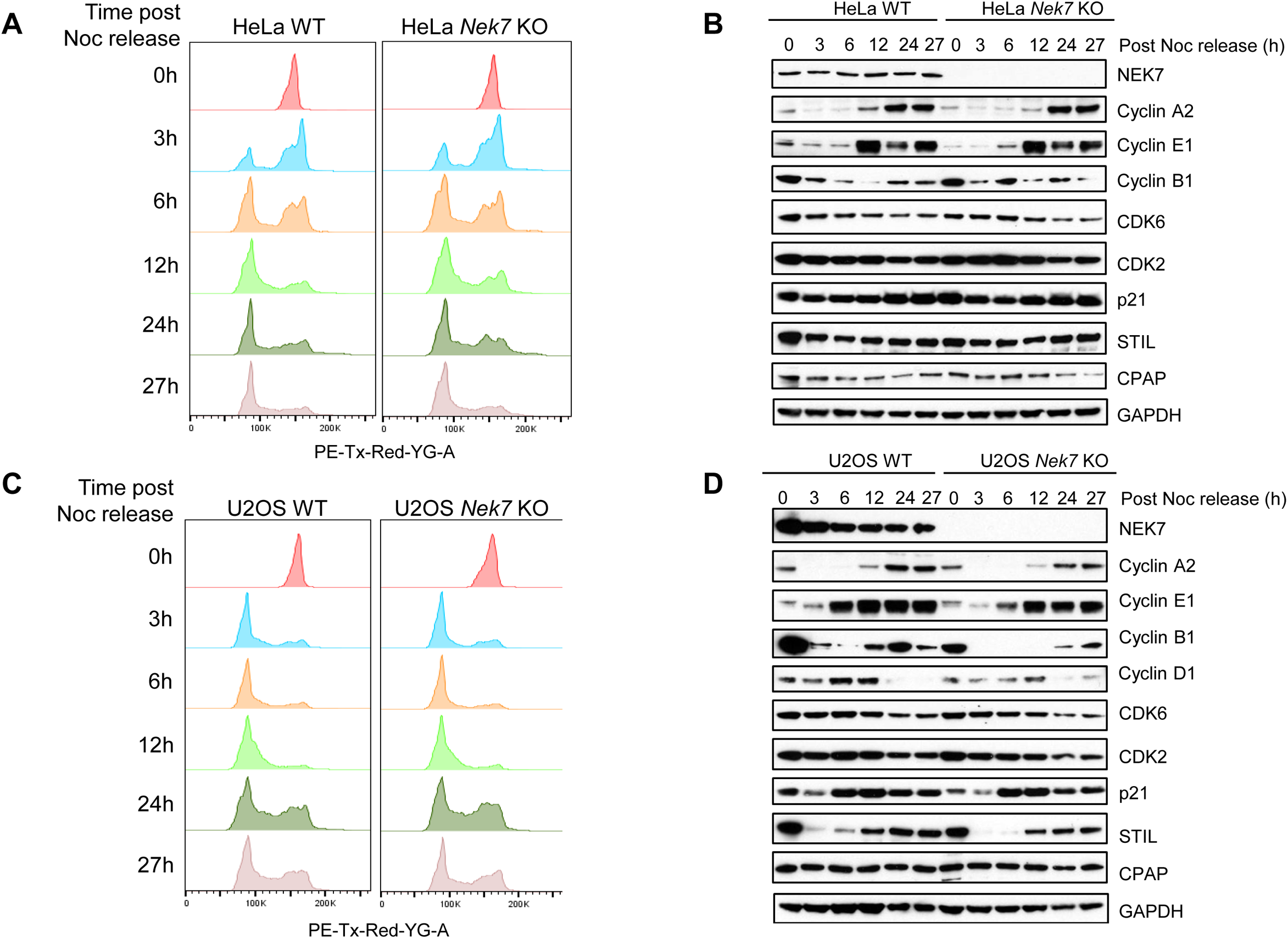
(A) WT and Nek7 KO HeLa cells were synchronized at early mitosis with 100 ng/mL nocodazole (Noc) for 16h. The cells were then released into fresh medium and collected at the indicated time points. The DNA content profiles for individual time points are shown. (B) Total cell lysates were analyzed by immunoblotting using antibodies against the indicated proteins. (C) WT and Nek7 KO U2OS cells were synchronized at early mitosis with 100 ng/mL nocodazole (Noc) for 16h. The cells were then released into fresh medium and collected at the indicated time points. The DNA content profiles for individual time points are shown. (D) Total cell lysates were analyzed by immunoblotting using antibodies against the indicated proteins.

**Supplementary Figure 7.**
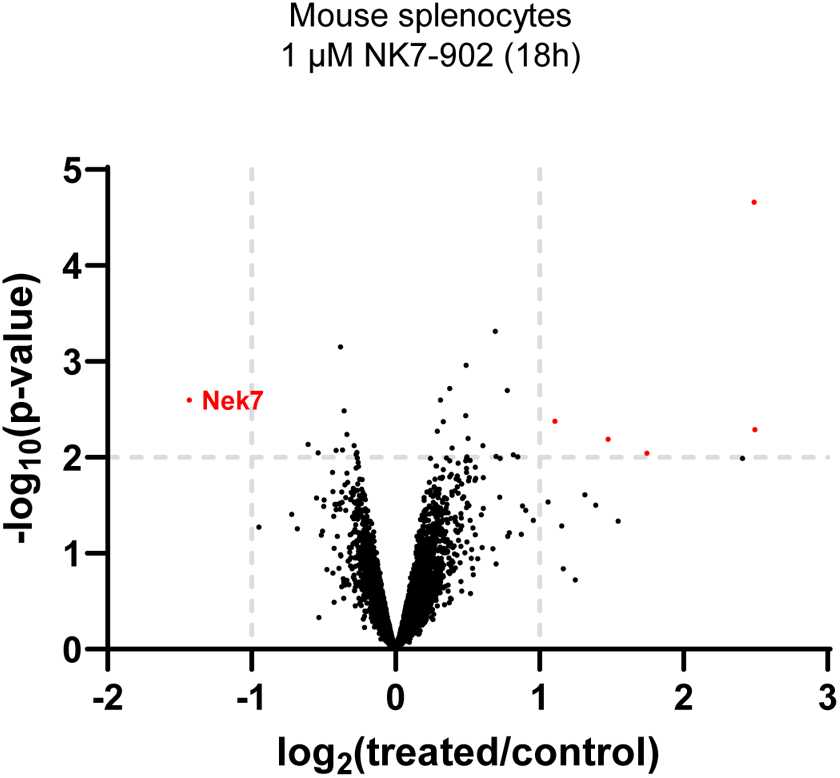
Mouse primary splenocytes were incubated with 1 µM NK7-902 for 18h and then analyzed by LC-MS/MS to identify differentially modulated proteins. Proteins in the top left quadrant were significantly downregulated (see also Supplementary Table 3).

**Supplementary Figure 8.**
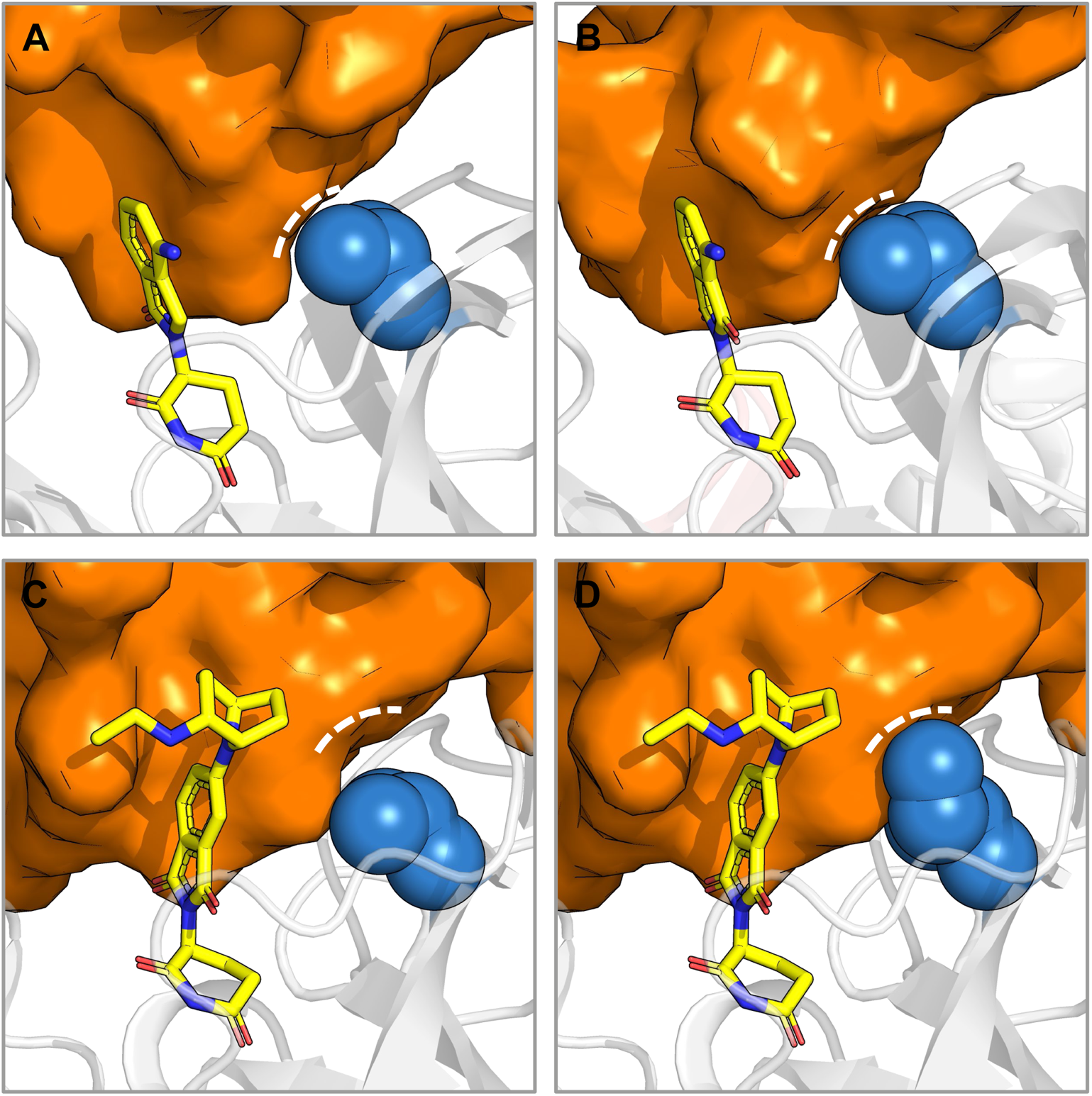
The crystal structures of (A) CRBN-lenalidomide-CK1α (5FQD) ^27^ and (B) CRBN-pomalidomide-IKZF1 (6H0F) ^31^ indicate that V388 of CRBN is tightly packed at the CRBN-protein of interest (POI) protein-protein interface. (C) The cryo-EM structure of CRBN-NK7-902-NEK7 indicates that V388 is more solvent exposed than in the CK1α or IKZF1 complexes. (D) Modelling the V388I point mutation indicates that isoleucine may be accommodated at the CRBN-NEK7 protein-protein interface without a steric clash. CRBN shown as grey cartoon, POI (CK1α / IKZF1 / NEK7) shown as orange surface, small molecule glue (lenalidomide / pomalidomide / NK7-902) shown as yellow sticks and residue 388 of CRBN shown as blue spheres.

**Supplementary Figure 9.**
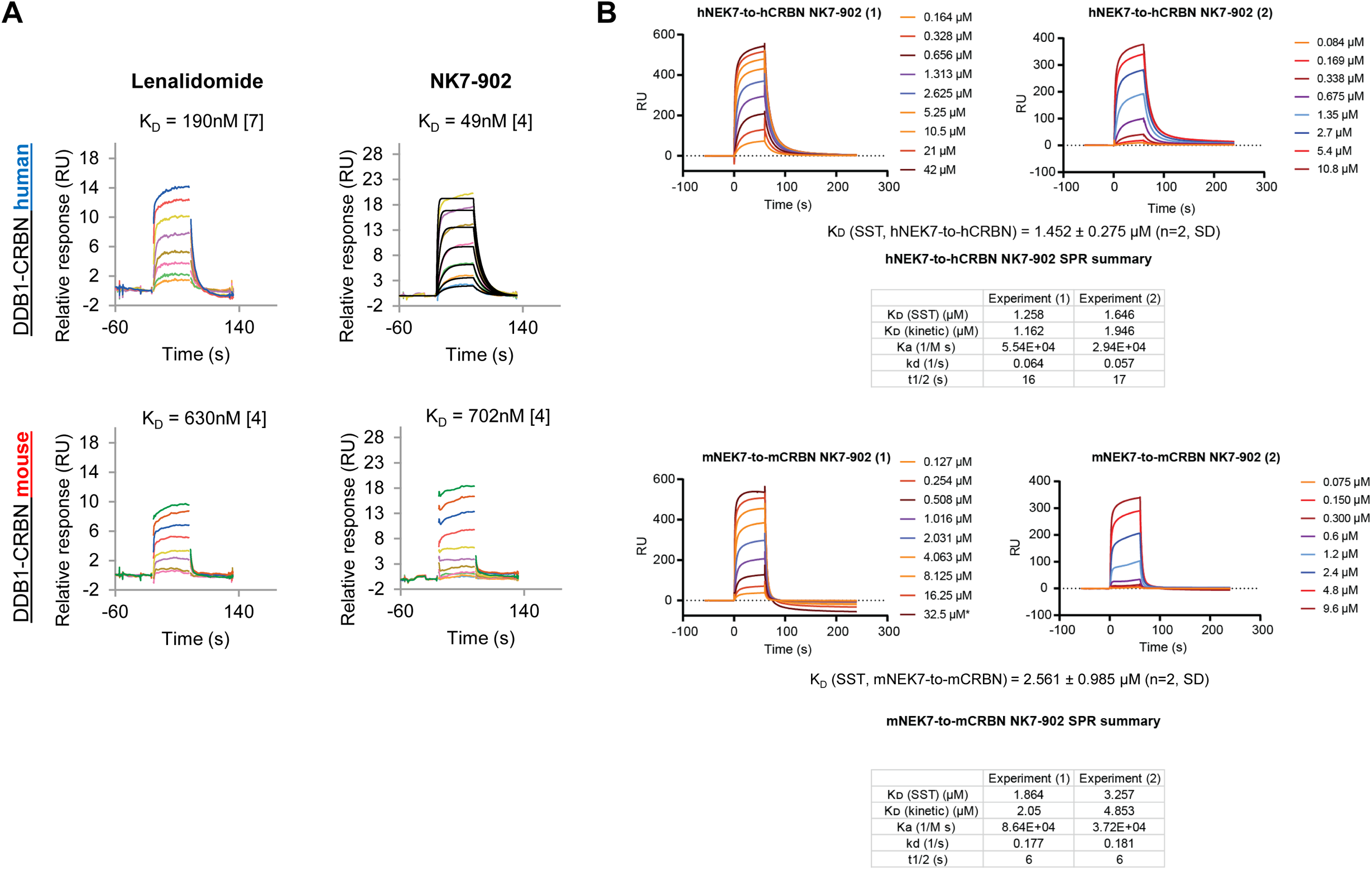
(A) SPR sensorgrams and KDs of the binary complex of both hCRBN/hDDB1 and mCRBN/mDDB1 to Lenalidomide and NK7-902. Displayed is the average of multiple (n) experiments described by the number in brackets. (B) SPR sensorgrams of the ternary complex of hNEK recruited to immobilized hCRBN//hDDB1 mediated by NK7-902. Shown are different concentrations of mNEK7 injected with a constant concentration of 5 µM NK7-902. Displayed below are steady state – derived fits of ternary complex formation by SPR of mNEK to mCRBN and hNEK to hCRBN of one representative experiment. (C) SPR sensorgrams of the ternary complex of hNEK recruited to immobilized mCRBN//mDDB1 and mNEK7 to hCRBN/hDDB1, both mediated by NK7-902. Shown are different concentrations of mNEK7/hNEK7 injected with a constant concentration of 5 µM NK7-902. The inlets display the steady-state derived fits of the respective SPR measurements.

**Supplementary Figure 10.**
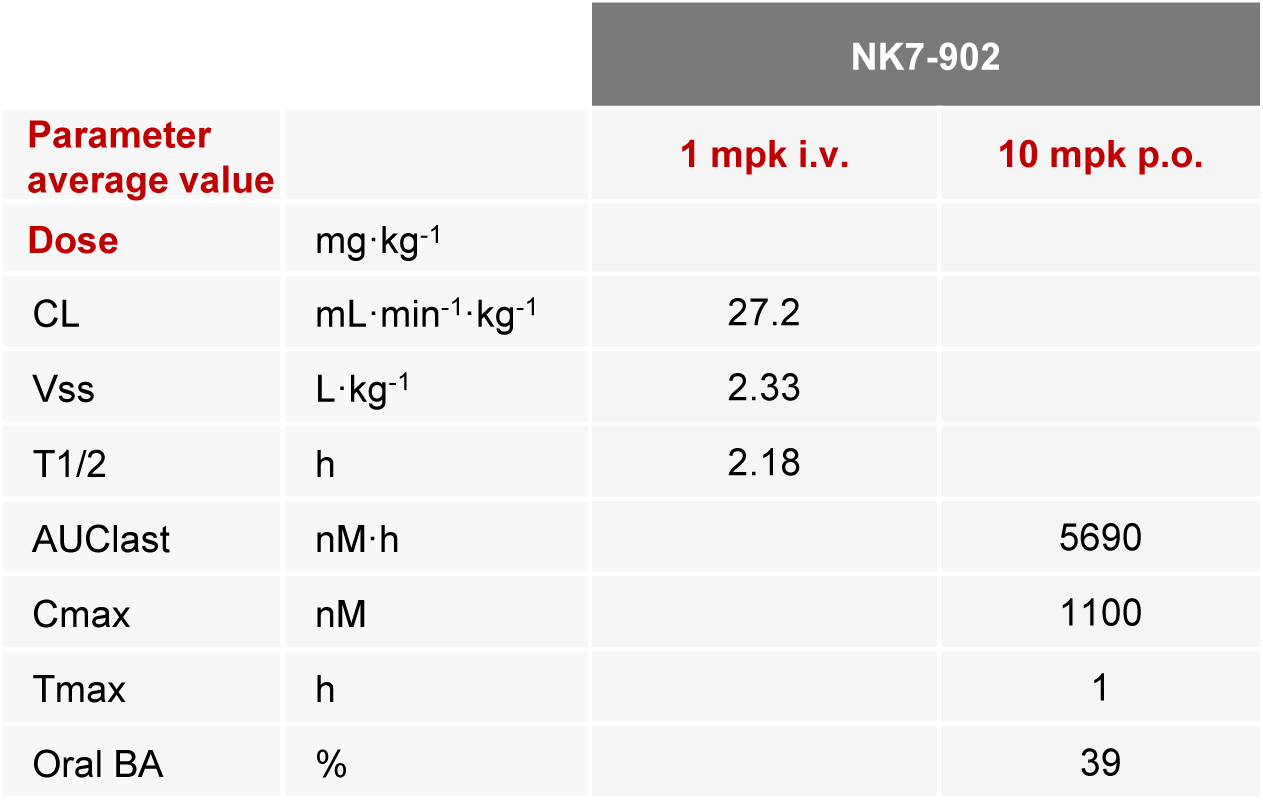
Mouse blood PK at the indicated doses.

**Supplementary Figure 11.**
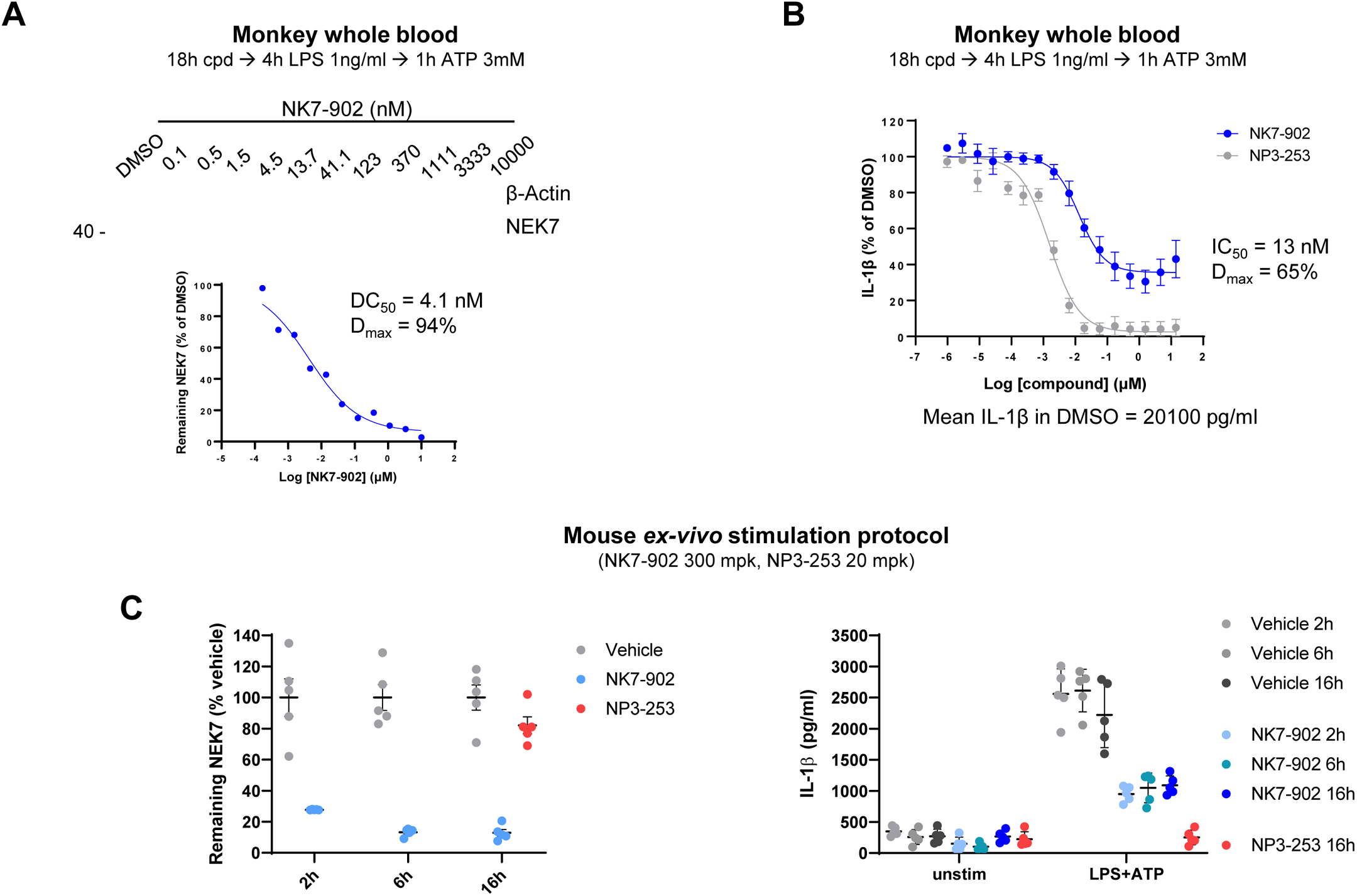
(A) NHP whole blood was diluted 1:2 in medium and incubated with increasing concentrations of NK7-902 for 18h prior to stimulation with LPS followed by ATP. Red blood cells were lysed and NEK7 and β-actin levels were measured in white blood cell protein extracts by Simple Western. Blood from four different monkeys was tested in separate experiments and a representative result is shown. (B) NHP whole blood was diluted 1:2 in medium and incubated with increasing concentrations of either NK7-902 or NP3-253 for 18h prior to stimulation with LPS and ATP. IL-1β levels in the plasma fraction were measured by HTRF. The mean IL-1β concentration (pg/mL) in the DMSO control is indicated below the graph. Data represent mean ± SEM (n=4). (C) Mice were dosed orally with either vehicle, 300 mg/kg NK7-902 or 20 mg/kg NP3-253 and blood samples were collected at the indicated times. Red blood cells were lysed and NEK7 and β-actin levels were measured in while blood cell protein lysates by Simple Western (left graph). At the same time points, blood samples were diluted 1:2 in medium and stimulated *ex-vivo* with 1 µg/mL crude LPS for 4h followed by an additional 45min with 3 mM ATP. IL-1β levels in the plasma fraction were measured by HTRF (right graph).

**Supplementary Figure 12.**
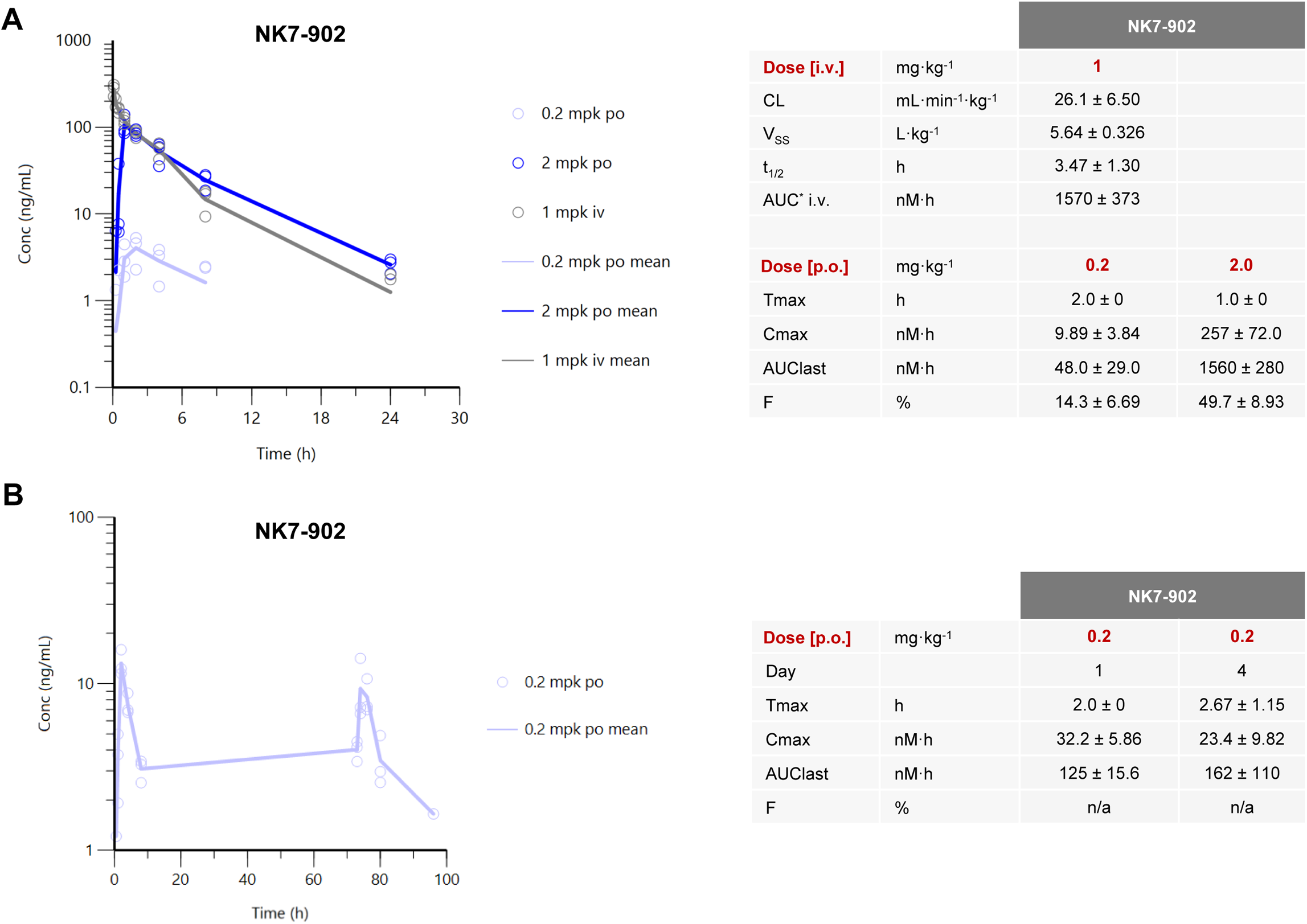
(A) Blood PK data for the single dose NHP PK/PD experiment shown in Figure 5A and 5B. (B) Blood PK data for the multiple-dose NHP PK/PD experiment show in Figure 5C.

**Supplementary Table 1.** Complete expression proteomics data in human primary monocytes treated with 1 µM NK7-902 for 18h.

**Supplementary Table 2.** Complete expression proteomics data in human iPS cells treated with 1 µM NK7-902 for 18h.

**Supplementary Table 3.** Complete expression proteomics data in mouse primary splenocytes treated with 1 µM NK7-902 for 18h.

